# Stomatal movement in *Arabidopsis* is driven by guard cell-localized and copper-insensitive CSD1 splice variant

**DOI:** 10.64898/2026.07.10.737675

**Authors:** Maryna Tsinyk, Kateřina Hlaváčková, Miroslav Ovečka, Jan Řehák, Jiří Sojka, Martina Špundová, Zuzana Kučerová, Jozef Šamaj, Tomáš Takáč, Petr Dvořák

**Affiliations:** Department of Biotechnology, Faculty of Science, Palacký University Olomouc, Olomouc, Czech Republic; Department of Biophysics, Faculty of Science, Palacký University Olomouc, Olomouc, Czech Republic

**Keywords:** superoxide dismutase, CSD1, FSD1, copper, SPL7, miR398, ABA-induced stomatal closure, reactive oxygen species, splice variant, guard cell

## Abstract

Copper (Cu) is an essential micronutrient whose bioavailability is strongly affected by soil physicochemical properties. During evolution, plants have developed mechanisms to flexibly adjust their metabolism to Cu status. Superoxide dismutases (SODs), including Cu/ZnSOD1 (CSD1) and FeSOD1 (FSD1), are key antioxidant enzymes regulated in Cu dependent manner in *Arabidopsis thaliana*. Examination of CSD1 cellular distribution and activity revealed that CSD1 is a nuclear and cytosolic SOD whose abundance and activity respond to Cu availability inversely to FSD1. Combined microscopic and biochemical analyses of Cu-dependent dynamics revealed that, unlike FSD1, CSD1 localization in guard cells (GCs) remains independent of Cu availability. *CSD1* escapes miR398-mediated regulation in GCs through a cell type-specific splice variant (*CSD1.2*) that carries an altered miR398-binding site. *In silico* analyses indicate that this mechanism is also present in crop species. Functionally, the *csd1* mutant showed reduced sensitivity to abscisic acid (ABA)-induced stomatal closure, a phenotype rescued by reintroducing *CSD1*. Biochemical and reactive oxygen species (ROS) level analyses indicate that CSD1.2 most likely acts independently of its canonical enzymatic activity in GCs and functions upstream of the ROS burst in the ABA signaling pathway. Together, we present a novel, cell-type-specific mechanism that safeguards ABA-driven stomatal closure under fluctuating Cu supply.

## Introduction

The early lifeforms evolved under anaerobic conditions in an ancient ocean saturated with redox-active transition metals such as iron (Fe), manganese (Mn), molybdenum (Mo), vanadium (V), chromium (Cr), and nickel (Ni) (Ivanovic and Vlaski-Lafarge, 2016). Among these, Fe was the most abundant (Wade et al., 2021) and predominantly existed as Fe^2+^, which served as the principal electron donor for anoxygenic photosynthesis (Olejarz et al., 2021). During the Great Oxygenation Event (GOE), the atmosphere became rich in oxygen (O_2_), leading to a drop in Fe bioavailability (David and Alm, 2011). In contrast, other essential metals, such as copper (Cu), became more soluble under aerobic conditions through the oxidation of Cu^1+^ to Cu^2+^ (Inupakutika et al., 2016). Both metals in trace amounts are essential for fundamental cellular processes. The strong redox activity of both metals enables rapid electron exchange, and this redox versatility is exploited in nature, as Cu- and Fe-containing enzymes drive key plant physiological processes such as photosynthesis or respiration (Ivanovic and Vlaski-Lafarge, 2016).

Aside from changes in metal deposition, an increase in O_2_ concentration led to the production of diverse reactive oxygen species (ROS) (Inupakutika et al., 2016), increasing the need for an effective antioxidant defense system in organisms. The diversification of the antioxidant defense system could be demonstrated by the evolution of superoxide dismutases (SODs), which transitioned from the ancient Fe/Mn-dependent isoforms to the later evolved Cu-dependent variants (Dreyer and Schippers, 2019).

SODs are characterized by high specificity for dismutation of superoxide radicals (O_2_^•–^) into hydrogen peroxide (H_2_O_2_) and O_2_ (Sheng et al., 2014). Considering substantial evidence for the signaling role of H_2_O_2_, SODs have strong potential to be considered among the driving forces in ROS-mediated signaling (Inupakutika et al., 2016; Řehák et al., 2026). Plant SODs are divided into FeSOD, Cu/ZnSOD, and MnSOD subfamilies based on their cofactor. The *Arabidopsis thaliana* genome encodes three isoforms of Cu/ZnSODs: cytosolic Cu/ZnSOD1 (CSD1), chloroplastic Cu/ZnSOD2 (CSD2), and peroxisomal Cu/ZnSOD3 (CSD3) (Kliebenstein et al., 1998). CSDs are activated by *Arabidopsis* COPPER CHAPERONE FOR SUPEROXIDE DISMUTASE (CCS) by the insertion of Cu^+^ into their active site (Cohu et al., 2009). FeSOD1 (FSD1) exhibits triple localization in plastids, cytoplasm, and nucleus (Dvořák et al., 2021), whereas FSD2 and FSD3 are localized to the chloroplast (Myouga et al., 2008). In the chloroplast, FSDs are activated through the CHAPERONIN 20 (Kuo et al., 2013).

Following the GOE, plants evolved complex mechanisms to cope with the increased Cu availability. At the front line, the COPPER TRANSPORTER (COPT) family serves as the essential hub for both initial root uptake and subsequent inter-organ distribution of Cu (Puig, 2014). Intracellularly, Cu is highly compartmentalized, precluding the presence of free ionic Cu in the cytosol (Burkhead et al., 2009). Chloroplasts and mitochondria serve as a primary Cu sink, while chloroplastic P-type ATPases of *Arabidopsis* (PAAs), such as PAA1 and PAA2, facilitate Cu import (Aguirre and Pilon, 2016). Cytosolic Cu is sequestered by specialized chaperones and metallothioneins, whereas the vacuole and cell wall serve as critical sites for sequestration, allowing the cell to buffer fluctuations in Cu availability (Burkhead et al., 2009; Klaumann et al., 2011). For example, the COPT5 transporter can transport Cu to the cytoplasm under Cu-deficient conditions (Klaumann et al., 2011; Puig, 2014).

In general, the response to Cu deficiency involves two main events: (i) activation of the Cu uptake system and (ii) redistribution of Cu among cuproproteins with subsequent down-regulation of non-essential ones (Yamasaki et al., 2007; Garcia-Molina et al., 2020). These processes are controlled by the transcription factor SQUAMOSA PROMOTER BINDING PROTEIN-LIKE7 (SPL7), which contains a recognition domain that binds to a GTAC core DNA motif in the promoters of target genes (Schulten et al., 2022). SPL7 induces expression of genes involved in Cu transport and storage, such as *COPT1, COP2,* and *COPT6*, or *FERRIC REDUCTASE OXIDASE 4* and *5* (*FRO4/FRO5*) (Sancenón et al., 2004; Seguí, 2017). SPL7 also regulates other cuproproteins by inducing several microRNA genes (*Cu-MIRs*), including *MIR397*, *MIR398*, *MIR408*, and *MIR857* (Perea-García et al., 2021). The main principle of the Cu economy is to conserve Cu for the most essential cuproproteins, such as plastocyanin (PC) and Cu oxidoreductases (COXs), which are crucial for photosynthesis (Burkhead et al., 2009). Instead, other cuproproteins are repressed by Cu-miRNAs, which often have multiple targets. CSDs and their chaperone, CCS, are also repressed by miR398 under Cu deficiency (Yamasaki et al., 2007; Beauclair et al., 2010). However, to replace downregulated CSDs, SPL7 directly induces *FSD1* expression, thereby enabling effective switching between SOD isoforms regardless of Cu availability (Yamasaki et al., 2009; Araki et al., 2018; Schulten et al., 2022). In Cu-rich conditions, SPL7-mediated regulation is attenuated by an unknown mechanism, leading to increased abundance and activity of CSDs (Garcia-Molina et al., 2020; Perea-García et al., 2021).

SOD isoform transition was shown to be crucial for chloroplast protection during oxidative stress induced by methyl viologen (MV). Phenotypic defects of the single *fsd1* mutant, under Cu deficiency (0.01 μM Cu), were restored to the wild-type levels by Cu supplementation (2 μM Cu) (Melicher et al., 2022). The protective functions are most likely accomplished by a chloroplast CSD2 isoform. A similar alteration can be proposed during salt-induced oxidative stress, as both FSD1 and CSD2 are associated with salt stress response (Dvořák et al., 2021; Zhuang et al., 2021; Melicher et al., 2026). FSD1 and CSD1 may also compensate for each other under contrasting Cu availability, since they share cytosolic localization. However, given their structural differences, it remains unclear to what extent CSD1/CSD2 and FSD1 are functionally redundant. To date, no visible developmental phenotype has been reported in *Arabidopsis* lacking CSD1. Nevertheless, the *csd1* mutant was found to be more heat-tolerant (Guan et al., 2013). Inhibition of CSDs by chemical inhibitor LUNG CANCER SCREEN 1 (LCS-1) caused oxidative stress and severe growth retardation in *Arabidopsis* (Frohn et al., 2024). Regarding other stress stimuli, *CSD1* expression was increased during plant responses to salt stress (Jagadeeswaran et al., 2009; Jia et al., 2009; Attia et al., 2011), light-induced stress (Xing et al., 2013), and pathogen infection (Jagadeeswaran et al., 2009). However, a cell-type-specific localization and function of CSD1 has yet to be reported.

This study aims to reveal the developmental expression and multi-level localization of CSD1-GFP, and to monitor how CSD1 and FSD1 respond to fluctuating Cu concentrations by tracking real-time changes in their abundance, activity, and distribution in complemented lines. Notably, we identified a stomatal guard cell (GC)-specific CSD1-GFP signal that remains stable regardless of Cu levels, whereas FSD1-GFP in GCs exhibits the usual Cu-dependent regulation. We propose that CSD1 participates in ABA-mediated stomatal closure, and the presence of a Cu-insensitive CSD1-GFP signal in GCs is explained by the circumvention of SPL7-miR398 regulation through alternative splicing of the miR398*-*binding site in the *CSD1* promoter. Finally, we have found that the role of CSD1 in ABA-induced stomatal closure is independent of its enzymatic activity and functions upstream of ABA-induced ROS production.

## Results

### Tissue-specific and subcellular localization of CSD1

CSD1 is considered to be a cytosolic protein (Kliebenstein et al., 1998); however, its subcellular, tissue-specific, and developmental localization has not been fully characterized.

To address this, we generated stable transformed lines expressing *CSD1* under the control of a native promoter (*pCSD1::CSD1:GFP::3′UTR-CSD1)* in the *csd1* mutant background (hereafter designated as CSD1-GFP line). The expression of the transgene and SOD enzymatic activity were validated by immunoblotting (Fig. S1A) and in-gel activity assays (Fig. S1B, C), respectively, and seedlings were used for subsequent advanced microscopy analyses.

The CSD1-GFP fluorescence signal remained consistently distributed across early developmental stages and was detected throughout the *Arabidopsis* seedling, with the highest intensity in the primary root tip (Fig. S2A-D). The signal was stable during the early stages of lateral root development (Fig. S2E-G) and showed enhancement in the transition and elongation zones of lateral roots at 7 days after germination (DAG) (Fig. S2G). CSD1-GFP line showed a signal concentration in the tips of growing root hairs (Fig. S2H). The aboveground tissues exhibited a homogeneous diffuse distribution at 4 DAG (Fig. S2I), while the signal intensity was generally lower than that of the root system (Fig. S2D).

A detailed Airyscan confocal laser-scanning microscopy revealed that the CSD1-GFP fusion protein localized to the cytoplasm and nuclei, but not to the nucleolus (Fig. 1A, B). The predominant nuclear enrichment was consistent in all aboveground tissues, including hypocotyl epidermis (Fig. 1A) and first true leaf epidermis (Fig. 1B). Line-profile analysis of fluorescence intensity revealed pronounced signal peaks corresponding to the nuclei, whereas signal intensity was markedly reduced in the nucleoli (Fig. 1G). In cotyledons, CSD1-GFP fluorescence was observed at various stages of stomatal development (Fig. 1C-F). Likewise, intensity profiling of mature stomata revealed two distinct peaks corresponding to the GC nuclei (Fig. 1H).

**Figure 1.**
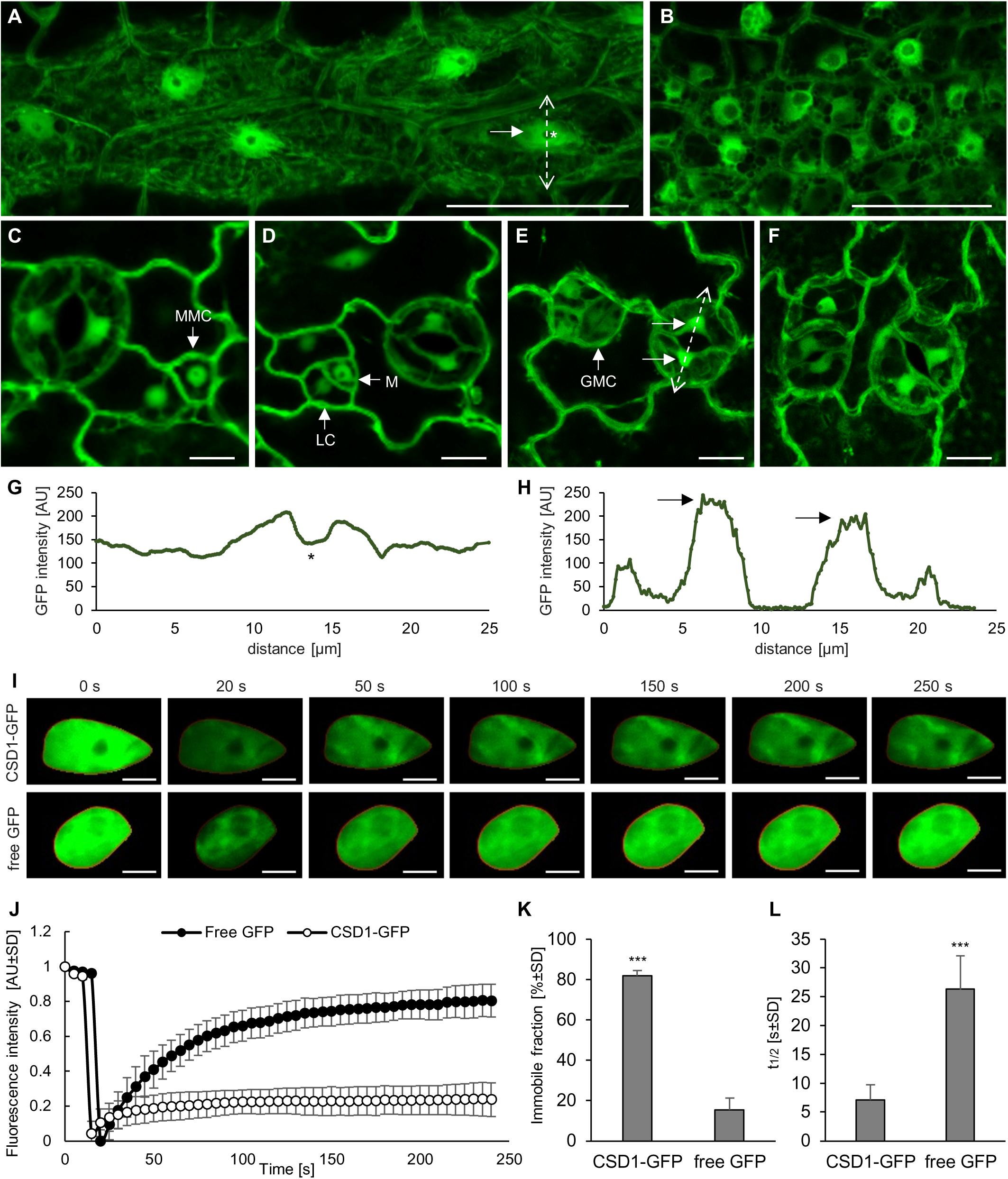
Subcellular localization and nuclear mobility analysis of CSD1-GFP in *Arabidopsis* cells. **(A)** Cells of hypocotyl epidermis with a strong nuclear (arrowhead) signal and no nucleolar (star) signal. **(B)** First true leaf epidermis. **(C-F)** CSD1-GFP signal within stomata lineage with (C) meristemoid mother cell (MMC), (D) meristemoid (M) and larger sister cell (LC), (E) dividing guard mother cells (GMC), and (F) mature stomata with a strong nuclear (arrowhead) signal. **(G, H)** Profile-based measurement of the fluorescence intensity distribution in (G) a hypocotyl epidermal cell and (H) mature stomata along the line shown in (A) and (E), respectively. Star correspond to labeled nucleolus in (A) and arrows to the labeled nuclei in (E). **(I-L)** Fluorescence recovery after photobleaching (FRAP) analysis of CSD1-GFP mobility in nuclei of hypocotyl epidermal cells. A transgenic line producing free GFP was used as a control. **(I)** Representative images of CSD1-GFP and free GFP nuclei before and after 250 s of selective bleaching. Time 0 s represents fluorescence intensity before bleaching, time 20 s represents fluorescence intensity immediately after bleaching, and time points 50-250 s show recovery of the fluorescence. **(J)** Fluorescence signal recovery in the examined nuclei of both lines. Fluorescence intensity values are shown in arbitrary units (AU) after normalization to the range from the highest intensities before bleaching (corresponding to 1) to the lowest intensities recorded after bleaching (corresponding to 0). **(K)** Relative proportion of immobile protein fractions of CSD1-GFP and free GFP in nuclei of respective expressing lines. **(L)** Average half-time of signal recovery for CSD1-GFP and free GFP in nuclei of respective expressing lines. Measurement was performed in three repetitions with a total number of examined seedlings n = 20 and a total number of analyzed nuclei n = 45 (for free GFP) and n = 53 (for CSD1-GFP). Error bars represent standard deviation. Stars indicate statistically significant differences between lines according to Student’s t-test (***P < 0.01). Scale bars – (A, B) 20 μm, (C-F) 10 μm, (I) 5 μm.

The localization of CSD1-GFP in both the cytosol and the nucleus prompts investigation of its nuclear-cytoplasmic transport. The molecular weight of CSD1-GFP is approximately 42.5 kDa, which is below the threshold for passive diffusion through the nuclear pore. Therefore, we performed a fluorescence recovery after photobleaching (FRAP) analysis with the CSD1-GFP line, using the free GFP as a positive control for passive diffusion (Fig. 1I; Fig. S3). Results showed nearly 20% FRAP of CSD1-GFP, compared to 80% recovery for free GFP, leaving almost 80% of the nuclear pool of CSD1-GFP immobile (Fig. 1J-L). These results indicate that most of the nuclear CSD1-GFP does not undergo passive nuclear-cytoplasmic exchange and remains immobile within the nucleus.

### CSD1 alternate FSD1 in sensing Cu fluctuations and holds Cu-independent pool in guard cells

As noted above, the abundances of FSD1, CSDs, and CCS depend on the Cu availability (Yamasaki et al., 2009). However, the kinetics of the Cu-dependent switch between CSD1 and FSD1 remain unknown, even though the speed of this transition can significantly influence the plant’s capacity to respond to stress.

Thus, microscopic analyses were conducted on complemented transgenic lines expressing GFP-fused SOD isoforms, which were transferred from conditions of sufficient Cu availability (2 μM Cu) to low Cu availability (0.01 μM Cu), and *vice versa* (Fig. 2A; Fig. S4). To minimize mechanical impact and maintain experimental consistency, hypocotyls of intact seedlings grown in Petri dishes on solid medium were monitored before and over 1-3 days after transfer (DAT) using a stereomicroscope. Fluorescence intensity was quantified across the entire hypocotyl region (Fig. S4). In parallel, to assess subcellular changes in signal distribution, cotyledons were imaged using CLSM (Fig. 2A). Additionally, CSD1, CSD2, CCS, and FSD1 abundance and SODs activity were monitored in transferred Col-0 seedlings, which were treated identically to the transgenic lines.

**Figure 2.**
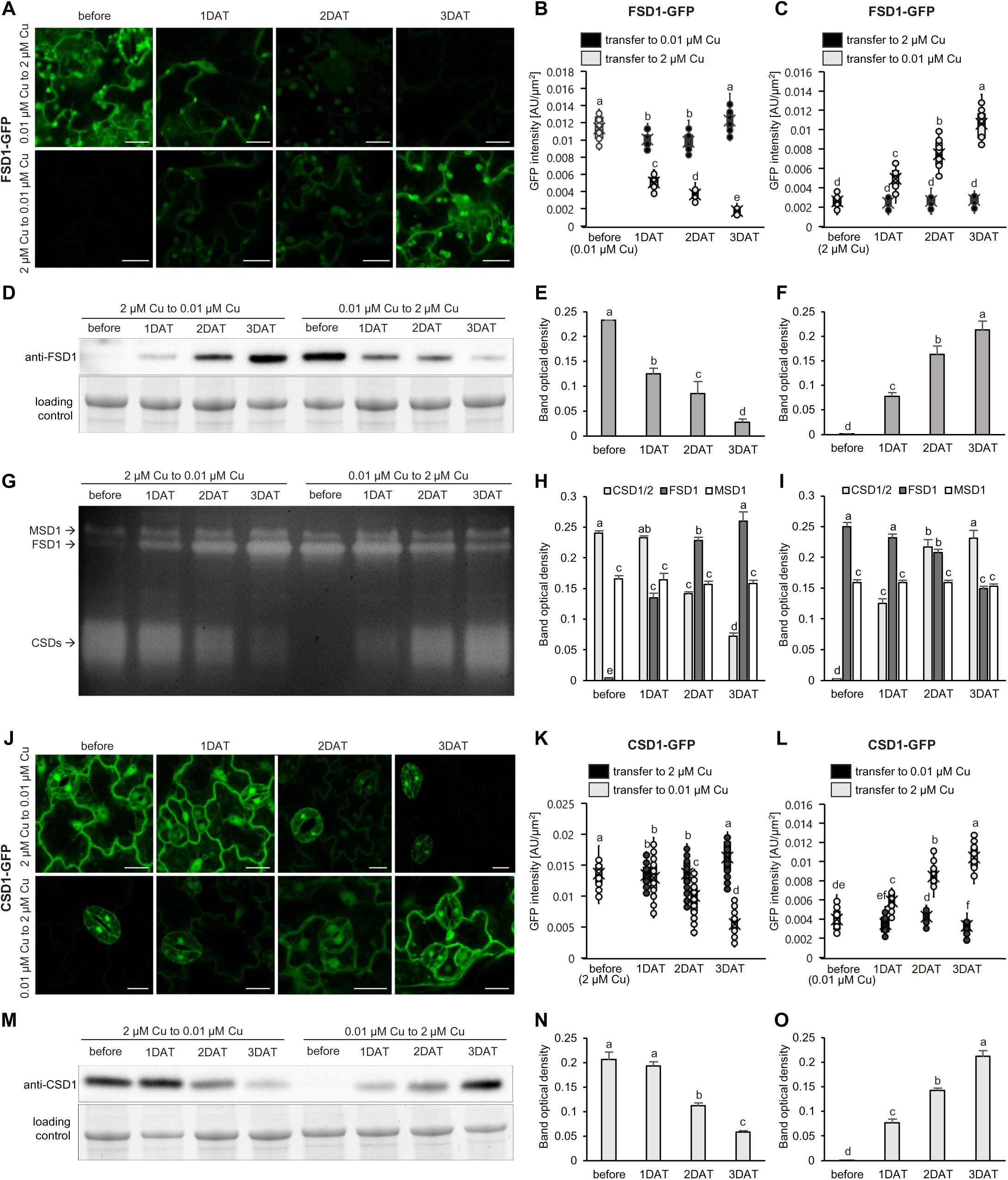
FSD1 and CSD1 response to changes in copper availability. **(A, J)** Representative images of the (A) FSD1-GFP and (J) CSD1-GFP cotyledons before and during 1-3DAT from 2 μM Cu to 0.01 μM Cu and from 0.01 μM Cu to 2 μM Cu. **(B, C, K, L)** Changes in fluorescence intensity in the (B, C) FSD1-GFP and in the (K, L) CSD1-GFP lines before and during 1-3 DAT (B, L) from 0.01 μM Cu to 2 μM Cu, where transfer from 0.01 μM Cu to 0.01 μM Cu was used as a control; and (C, K) from 2 μM Cu to 0.01 μM Cu, where transfer from 2 μM Cu to 2 μM Cu was used as a control. Values are calculated as arbitrary units (AU) per measured hypocotyl area (μm²). The experiment was performed in triplicate, with a total of n = 50 examined seedlings. Levels not connected by the same letter differ significantly (one-way ANOVA, P<0.01). **(D-F, M-O)** Abundance of (D) FSD1 and (M) CSD1 in Col-0 seedlings before and during 1- 3 DAT from 2 μM to 0.01 μM Cu and from 0.01 μM to 2 μM Cu, with statistical evaluation of changes during transfer (E, O) from 0.01 μM to 2 μM Cu and (F, N) from 2 μM to 0.01 μM Cu. **(G)** Visualization of endogenous SOD isozyme activities on native polyacrylamide gel and (H, I) statistical evaluation of isoenzyme activity before and during 1-3 DAT in seedlings (H) from 2 μM to 0.01 μM Cu and (I) from 0.01 μM to 2 μM Cu. The experiment was performed in triplicate, with a total of n = 90 examined seedlings. Error bars represent standard deviation. Levels not connected by the same letter differ significantly (one-way ANOVA, P<0.01). Scale bar – 20 μm.

Under Cu-deficient conditions, FSD1-GFP was localized to the cytoplasm, chloroplasts, and nuclei, with an initial fluorescence signal intensity of 0.011 unit/µm^2^ (Fig. 2A, B). Upon transfer to 2 μM Cu, the signal intensity decreased by nearly half at 1 DAT and continued to decline to 0.15-fold of the initial level at 3 DAT (Fig. 2A, B). Interestingly, immunoblotting of endogenous FSD1 confirmed this depletion trend, although FSD1 activity remained stable at 1 DAT and began to decline in subsequent days (Fig. 2D, E, G, I). Moreover, we employed *in vivo* light-sheet fluorescence microscopy (LSFM) for time-lapse monitoring of hypocotyl epidermal cells and precisely determined the timing of the decrease in FSD1-GFP fluorescence intensity. This microscope modality maintains seedlings under near-physiological conditions during long-term experiments by providing light access to the green part and automated media exchange via a peristaltic pump to roots, effectively mimicking natural growth environments. The observation revealed that the FSD1-GFP signal began to decline approximately 20 h after the media exchange (Video S1 and S2), consistent with the results described above. Conversely, transferring the FSD1-GFP line from 2 to 0.01 μM Cu resulted in a gradual restoration of fluorescence. Intensity increased nearly twofold at 1 DAT and reached 0.0106 unit/µm^2^ by 3 DAT (Fig. 2A, C). This recovery was mirrored by endogenous FSD1 protein and activity levels (Fig. 2D, F, G, H).

CSD1-GFP initially exhibited strong cytosolic and nuclear signals (0.014 units/µm^2^) under sufficient Cu supply (Fig. 2J, K). Following transfer to 0.01 μM Cu, the decrease in signal was less pronounced and remained unchanged after 1 DAT (Fig. 2J, K). By the 3DAT, total signal intensity reached 0.33-fold of the initial levels (Fig. 2J, K). Notably, while fluorescence was markedly reduced in pavement cells (PCs), a strong signal persisted in the cytoplasm and nuclei of stomatal GCs (Fig. 2J). Separate quantification at the cellular level revealed that PCs exhibited a 0.20- to 0.25-fold decline in CSD1-GFP signal intensity in the cytoplasm and nuclei, respectively, whereas the signal in GCs remained at 0.80- to 0.98-fold of its initial intensity (Fig. S5). This GC-specific persistence in seedlings grown on low Cu suggests a localized physiological requirement rather than residual Cu storage (Fig. 2J). In Col-0 wild-type plants, the abundance of CSD1, CSD2, and CCS remained stable at 1 DAT, indicating a delayed response to Cu deficiency (Fig. 2M, N; Fig. S6A, B, D, E). Total CSD activity correlated closely with protein abundance (Fig. 2G, H). LSFM imaging of CSD1-GFP showed a significant initial decrease approximately 30 h after the media were exchanged (Videos S3 and S4), which is consistent with the findings described above. Transfer back to Cu-sufficient media induced a steady increase in CSD1-GFP fluorescence intensity, reaching 0.0104 units/µm^2^ by 3 DAT (Fig. 2J, L). These results were consistent with the restoration of endogenous CSDs, CCS protein levels, and total CSD activity (Fig. 2G, I, M, O; Fig. S6A, C, D, F).

Seedlings transferred to media with identical Cu concentrations exhibited minor fluctuations in signal intensity but remained relatively stable (Fig. S7A, B). Moreover, no significant differences in either protein abundance (Fig. S8A) or SOD activity (Fig. S8B) were observed, thereby excluding mechanical stress effects.

Our results suggest that Cu fluctuations are sensed concurrently by FSD1 and CSD1 within 20-30 h, though they exhibit different reduction rates.

### csd1 mutants exhibit dysregulated stomatal closure in response to ABA

Our results raise a question about the role of CSD1 in the stomata. Therefore, we have examined the ABA-induced stomatal closure of both *csd1* mutant lines and the complemented CSD1-GFP line under Cu sufficiency. Mock treatment showed fully opened stomata in all treated lines (Fig. 3A, B). When seedlings were treated with 50 μM ABA, *csd1* mutants did not respond by stomatal closure, in contrast to the Col-0 and complemented CSD1-GFP line, which showed closed stomata (Fig. 3A, B). These observations indicate a CSD1 function in ABA-induced stomatal closure. The same experiment was conducted with seedlings grown under Cu deficiency to exclude the possible impact of high Cu concentration and FSD1 interference, and it showed similar results (Fig. S9A, B).

**Figure 3.**
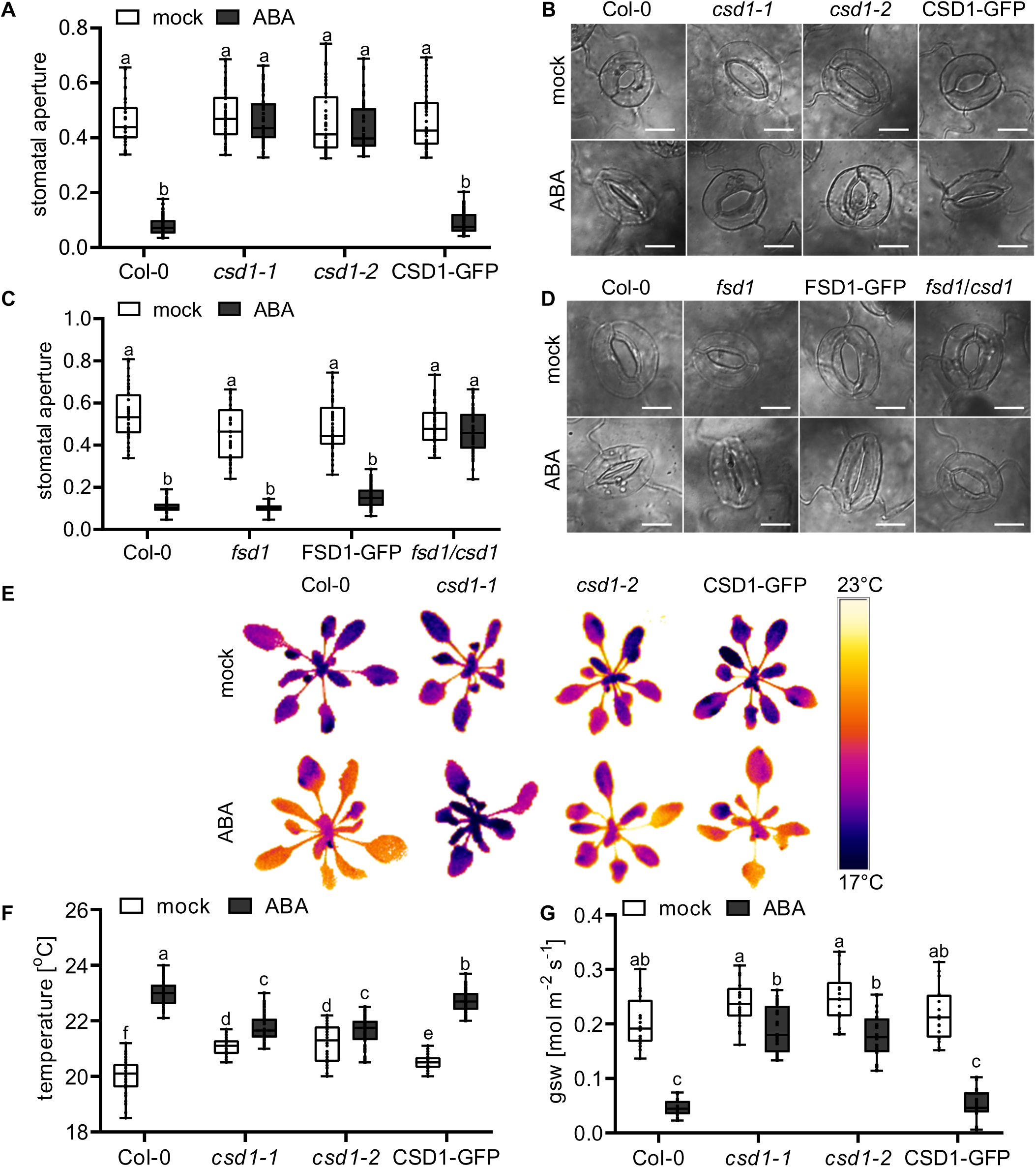
The *csd1* mutant displays dysregulated ABA-induced stomatal closure independent of Cu availability. Stomatal aperture of Col-0, single *csd1* and *fsd1*, and double *fsd1/csd1* mutants and complemented CSD1-GFP and FSD1-GFP lines was measured after application of 50 μM abscisic acid (ABA). **(A, C)** Quantification of stomatal aperture (ratio of stomata pore width to length) in 7-day-old seedlings grown on (A) 2 μM or (C) 0.01 μM Cu treated with opening buffer with or without ABA. The experiment was performed in triplicate (n = 50). **(B, D)** Representative images of stomata in 7-day-old seedlings grown on (B) 2 μM 1/2 MS or (D) 0.01 μM 1/2 MS treated with opening buffer with or without ABA. **(E)** Representative thermal images of rosettes from 4-week-old plants of Col-0, *csd1* mutants, and CSD1-GFP lines treated with or without 100 μM ABA. **(F)** Leaf temperature from 4-week-old plants of Col-0, *csd1* mutants, and the CSD1-GFP lines treated with or without 100 μM ABA. 20 temperature values from thermal images captured by a thermal camera were assessed per plant (n = 60). **(G)** Leaf stomatal conductance to water vapour (gsw) in 4-week-old plants of Col-0, *csd1* mutants, and CSD1-GFP lines treated with or without 100 μM ABA. The gsw was estimated from 50-80 leaves from 20 individual plants (n = 20). Levels not connected by the same letter differ significantly (one-way ANOVA, P<0.01). Scale bar – 10 μm.

It was also of interest to test ABA-induced stomatal movements in the *fsd1* mutant and newly-prepared *fsd1/csd1* double mutant to examine the Cu-dependent compensatory mechanism between CSD1 and FSD1. The *fsd1/csd1* double mutant was validated as a stable homozygous line (Fig. S10). Under Cu-deficient conditions, the stomata of *fsd1* and FSD1-GFP lines showed ABA sensitivity similar to Col-0 (Fig. 3C, D). In contrast, the stomata of *fsd1/csd1* double mutant remained open in response to ABA (Fig. 3D). Furthermore, these phenotypes were reproduced in seedlings grown under Cu-sufficient conditions (Fig. S9C, D). Together, these findings support a unique, Cu-independent role for CSD1 in ABA-mediated stomatal closure.

To confirm the impairment of stomatal closure observed in *csd1* mutants with independent methods, we measured changes in leaf temperature and stomatal conductance for water vapour (gsw) following ABA treatment in 4-week-old plants. Col-0 and the complemented CSD1-GFP line showed an increase in leaf temperature after ABA application (Fig. 3E, F). Due to ABA-induced stomatal closure and consequent reduction of leaf cooling by transpiration, leaf temperature increased by 2.9℃ and 2.2°C in Col-0 and CSD1-GFP, respectively (Fig. 3E, F). Instead, leaf temperature difference between mock and ABA-treated plants of *csd1-1* and *csd1-2* mutant lines was only 0.6℃ and 0.5℃, respectively (Fig. 3E, F). These results corresponded to the changes in gsw following ABA treatment, showing that *csd1-1* and *csd1-2* mutants exhibited a less pronounced decline, approximately 3-fold smaller than Col-0 and CSD1-GFP (Fig. 3G), thus confirming an impaired stomatal response to ABA treatment. Overall, the results suggest that CSD1 is required for stomatal closure in response to ABA.

### Alternative splicing confers CSD1 insensitivity to miR398

The presence of CSD1-GFP signal in stomata in Cu deficient conditions (Fig. 2J) can be explained by two different reasons: 1) either *MIR398* is not expressed in GCs, preventing the downregulation of *CSD1*; or 2) the SPL7-miR398-mediated regulation is disrupted. To answer the first question, we generated transgenic lines carrying a nuclear-localized *GFP* under the control of the native promoters of *MIR398a*, *MIR398b*, and *MIR398c*. All three lines, germinated and grown on low Cu, showed a GFP-specific fluorescent signal in the nuclei of both PCs and GCs, proving a simultaneous presence of CSD1 and its negative regulators in GCs (Fig. 4A). Instead, under sufficient Cu supply, the lines showed no detectable signal in the nuclei of PCs and GCs (Fig. S11). These data provided clear evidence about the Cu-dependent regulation of *MIR398s*.

**Figure 4.**
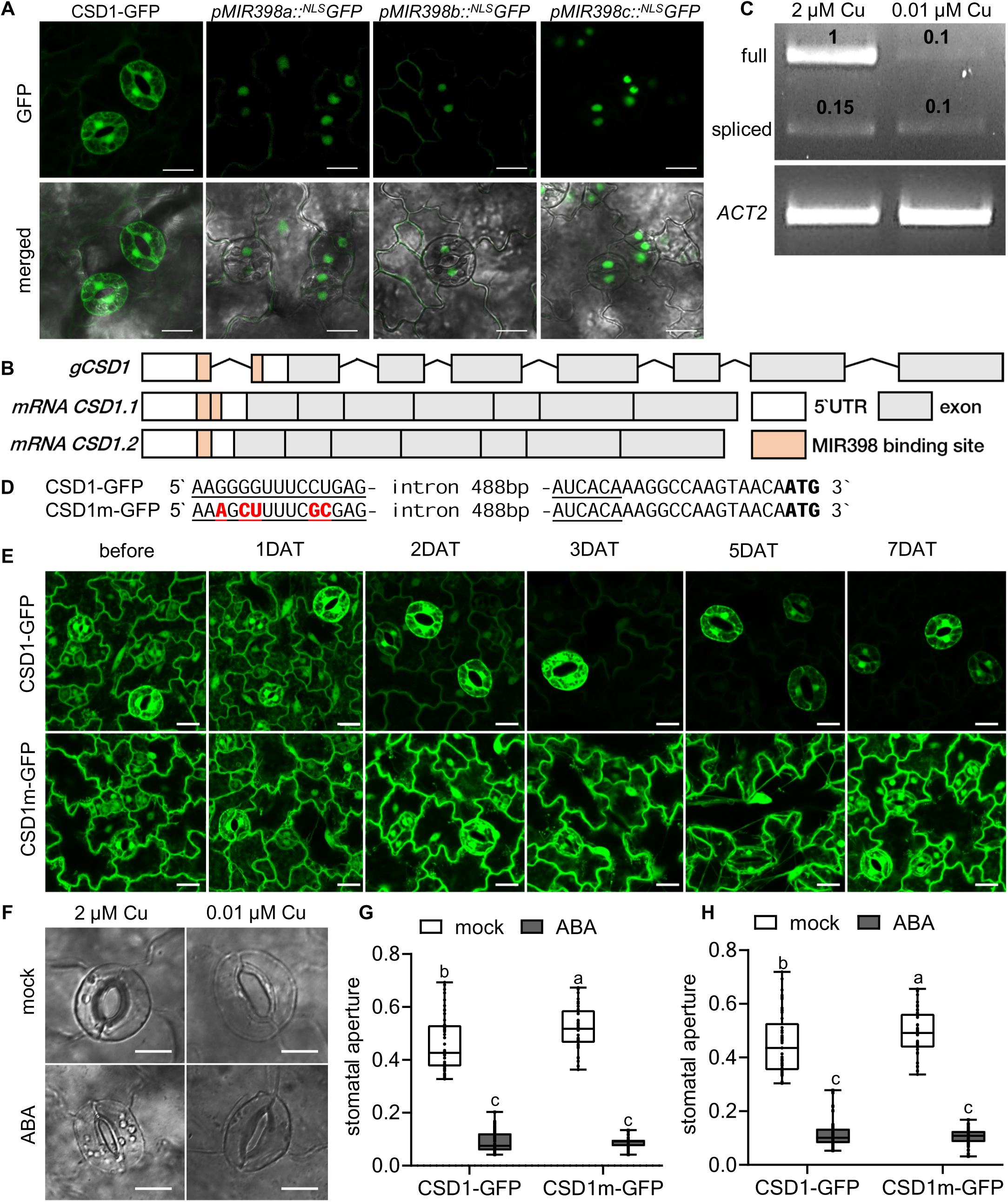
miR398 localization and Cu-insensitive variant of CSD1. **(A)** Localization of CSD1-GFP and nuclear-localized GFP (*^NLS^GFP*) driven by a *MIR398s* promoter under low Cu conditions (0.01 µM Cu). **(B)** Scheme of the *CSD1* gene and splicing variants sensitive (*CSD1.1*; NM_100757.4) and insensitive to miR398 (*CSD1.2*; NM_001084025.1). **(C)** Full (512 bp) and spliced (402 bp) variants were determined by reverse transcription PCR analysis in the *Arabidopsis* Col-0 seedlings grown on 2 µM Cu and 0.01 µM Cu with two forward primers (primer scheme in Figure S12B). Numbers indicate band intensity ratios relative to the full variant at 2 µM Cu (n = 3). *ACTIN2* is used as an internal control. **(D)** Fragment of the CSD1-GFP and CSD1m-GFP genomic sequences with an underlined miR398 binding site and highlighted nucleotides (red), which were mutated in the CSD1m-GFP line. **(E)** Representative images of CSD1-GFP and CSD1m-GFP seedlings before and during 1-7 days after transfer (DAT) from 2 μM Cu to 0.01 μM Cu. **(F)** Representative images of stomata in 7-day-old CSD1m-GFP seedlings grown on 2 μM Cu and 0.01 Cu treated with opening buffer with or without ABA (50 μM). **(G, H)** Quantification of stomatal aperture (ratio of stomata pore width to length) in 7-day-old seedlings grown on (G) 2 μM Cu or (H) 0.01 μM Cu treated with opening buffer with or without ABA (50 μM). The experiment was performed in triplicate, with a total of n = 50 stomata. Levels not connected by the same letter differ significantly (one-way ANOVA, P<0.01). Scale bars – (A, D) 20 μm, (F) 10 μm.

Such observation raises the question of how CSD1 bypasses Cu-SPL7-miR398 regulation, specifically in GCs. The miR398-specific binding site (BS) of *CSD1* is located in the 5′UTR, and it is disrupted by an intron (486 bp) (Fig. 4B). The normally spliced variant of *CSD1* links the two parts of the BS to allow the miR398 binding (*mRNA CSD1.1; NM_100757.4*) (Fig. 4B; Fig. S12A). Alternative splicing involves a 9-bp downstream shift of the 3′ splice site, resulting in the exclusion of the second part of the miR398 BS (*mRNA CSD1.2; NM_001084025.1*) and thus preventing the miR398 binding (Fig. 4B; Fig. S12A). We performed RT-PCR analysis to determine the abundance of *CSD1.1* and *CSD1.2* mRNAs in Col-0 seedlings under different Cu availability. The single reverse primer was used simultaneously with two forward primers. The first one attached to the start of the 5′UTR and the second one was designed to anneal within the spliced BS in *CSD1.2*, yielding 512 bp (*CSD1.1*) and 402 bp (*CSD1.2*) products, respectively (Fig. S12B). Consistent with our previous results, the full-length variant was the most abundant in seedlings grown on 2 μM Cu, approximately 10 times more than in seedlings grown on 0.01 μM Cu (Fig. 4C). Instead, the spliced variant showed similar abundance under both Cu-sufficient and Cu-deficient conditions (Fig. 4C), suggesting that the splicing event is Cu-independent. Thus, we demonstrate that under low-Cu availability, *CSD1* can bypass the miR398 module via alternative splicing within the miR398-specific BS.

To study the dependence of CSD1 on Cu and its main regulators, miR398s, we introduced 5 point mutations in the miR398-specific BS of *CSD1* (Fig, 4D), generating a potentially miR398-insensitive line in the *csd1-1* background (hereafter referred to as the CSD1m-GFP line). The CSD1m-GFP line grown on 2 μM Cu showed a fluorescence signal pattern similar to the complemented CSD1-GFP line, with a CSD1 localization in the cytoplasm and nuclei of cotyledon epidermal cells (Fig. 4E). In contrast to CSD1-GFP, the CSD1m-GFP fluorescent signal remained stable even after 7 DAT on 0.01 μM Cu (Fig. 4E; Fig. S13A), confirming that miR398 can be considered the principal Cu-dependent regulator of CSD1. Consistent with the microscopic analysis, immunoblotting with anti-CSD1 antibody showed that the protein level in the CSD1m-GFP line remained stable (Fig. S13B). The abundance of CCS (essential for proper CSD1 activity) decreased in both Cu-sensitive and Cu-insensitive lines, following a pattern similar to that observed in Col-0 (Fig. S13B). The reduction in CSD1 activity correlated with the decline in CCS levels in both lines, even though CSD1 abundance remained unchanged in the CSD1m-GFP line (Fig. S13C). Further, we wondered whether CSD1m-GFP could rescue the *csd1* mutant phenotype in ABA-induced stomatal closure. CSD1m-GFP seedlings showed a decrease in stomatal aperture after ABA treatment comparable to the CSD1-GFP line in both Cu sufficient and deficient conditions (Fig. 4F-H).

These results indicate that the persistent presence of CSD1 in GCs is attributable to cell-specific alternative splicing within the miR398 BS. Moreover, while miR398-resistant CSD1 maintains high abundance under low Cu conditions, its activity remains limited by CCS availability. As a result, the requirement of CSD1 for ABA-induced stomatal closure appears to be independent of its enzymatic activity.

### The guard cell pool of CSD1 is required for stomatal closure regardless of its activity

Although both CSD1-GFP and CSD1m-GFP lines successfully rescued the *csd1* mutant phenotype under all Cu conditions, it remained unclear whether the role of CSD1 in ABA-induced stomatal closure could be attributed solely to GC-specific CSD1. To address this, we prepared a transgenic line expressing CSD1-GFP under a GC-specific promoter (hereafter referred to as GC-CSD1-GFP line). The exchange of the promoter caused miR398-insensitivity due to the lack of miR398 BS (present in the 5′UTR sequence of the *CSD1* gene). As a result, the transgenic line showed a fluorescent signal in the stomata of seedlings grown on both 2 μM and 0.01 μM Cu (Fig. 5A). Immunoblotting with an anti-CSD1 antibody showed that the CSD1-GFP abundance in this line is stable across Cu concentrations (Fig. S14A), while the SPL7-miR398 regulatory module was present and functional, as indicated by intensive decrease in CSD2 abundance under low Cu conditions (Fig. S14A).

**Figure 5.**
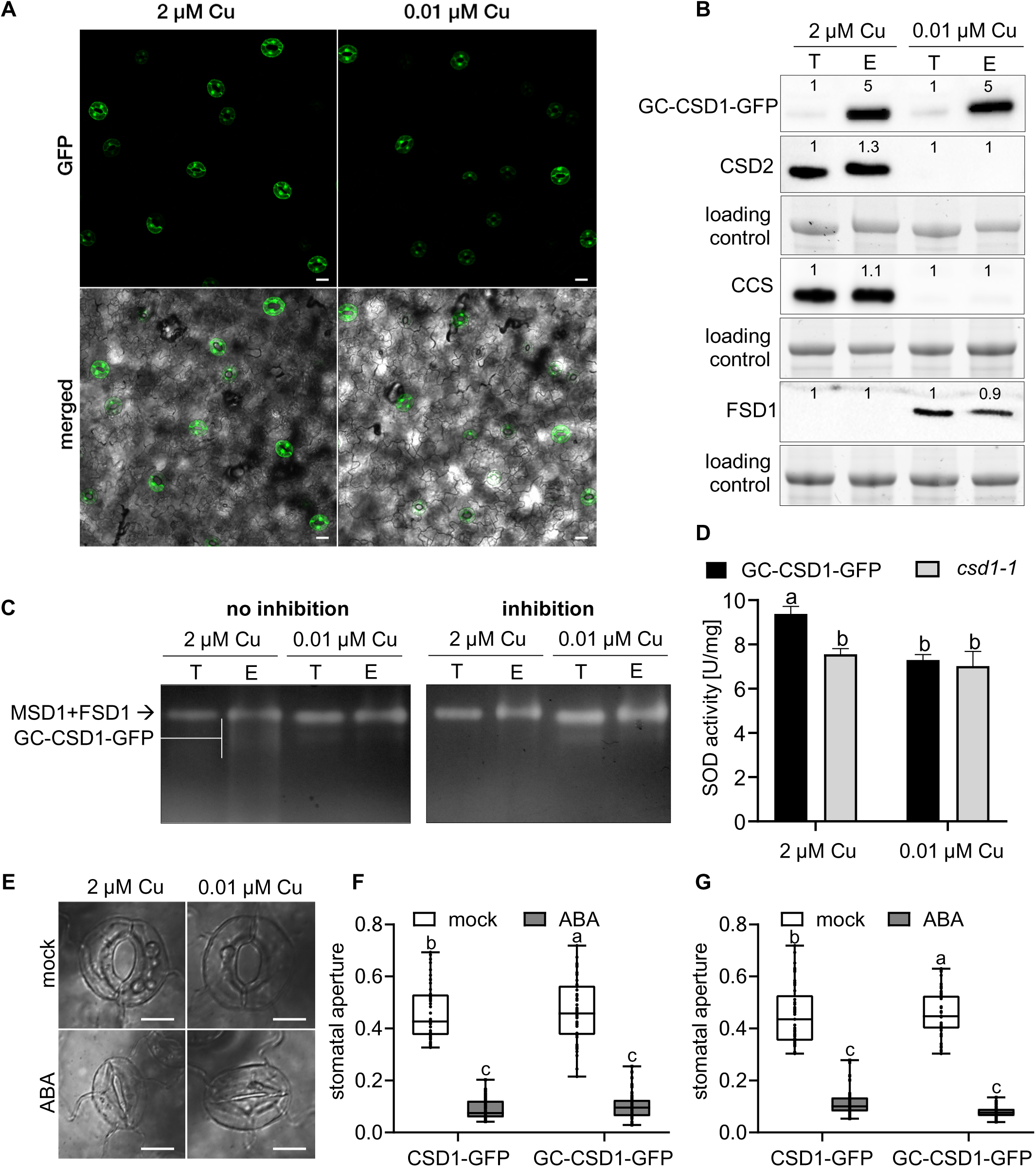
Characterization of the CSD1 guard cell pool. **(A)** Localization of CSD1-GFP under a guard-cell-specific promoter in7seedlings grown on 2 µM Cu and 0.01 µM Cu. **(B)** Protein abundance in the total extract (T) and the extract of guard-cell-enriched fraction (E) obtained from 14-day-old GC-CSD1-GFP seedlings grown on 2 µM Cu and 0.01 µM Cu. Numbers show the ratio of normalized band intensity (total/enriched) at each Cu concentration. **(C)** Visualization of SOD enzyme activity on native gel with and without inhibition by KCN (which inhibits CSD activity) in the total extract (T) and the extract of guard-cell-enriched fraction (E) obtained from 14-day-old GC-CSD1-GFP grown as described above. **(D)** Total SOD activity in the guard-cell-enriched extract of GC-CSD1-GFP and *csd1-1* lines grown as described above. **(E)** Representative images of stomata in 7-day-old GC-CSD1-GFP seedlings grown on 2 μM and 0.01 µM Cu treated with opening buffer with or without ABA (50 μM). **(F, G)** Quantification of stomatal aperture (ratio of stomata pore width to length) in 7-day-old seedlings grown on (F) 2 μM Cu or (G) 0.01 μM Cu treated with opening buffer with or without 50 μM ABA. The experiment was performed in triplicate, with a total of n = 50 stomata. Levels not connected by the same letter differ significantly (Student’s t-test, P<0.01 (D); one-way ANOVA, P<0.01 (F, G)). Scale bars – (A) 20 μm, (E) 10 μm.

To increase the pool of CSD1-GFP, we enriched GC fractions from the GC-CSD1-GFP seedlings grown on both 2 μM and 0.01 μM Cu (Fig. S14B). Enriched fractions showed a 5-fold increase in CSD1-GFP abundance relative to the total extract (Fig. 5B), confirming the Cu-independent, GC-specific expression of *CSD1* in this line. In contrast, CSD2, CCS (at high Cu) and FSD1 (at low Cu) pools remained almost unchanged after enrichment (Fig. 5B). CSD1-GFP activity in this line was almost undetectable in the total extract (Fig. 5C). However, in the enriched extract, the KCN-sensitive, CSD1-GFP-specific band was observed only in samples from plants grown under high Cu availability (Fig. 5C), indicating the inactivity of GC-specific CSD1 on low Cu. For CSD1 activity quantification, we also measured the total SOD activity in enriched extract spectrophotometrically. To exclude possible interference of other SOD isoform activities, we measured the total SOD activity in the enriched extract from the *csd1* mutant. Under low Cu conditions, the total activity in the *csd1* mutant and the GC-CSD1-GFP line was comparable (Fig. 5D). However, in extracts from plants grown on 2 µM Cu, the activity in the GC-CSD1-GFP line significantly increased (Fig. 5D), suggesting that the CSD1 activity in this line also occurs exclusively under Cu-sufficient conditions.

We also tested whether the GC-CSD1-GFP construct could rescue the *csd1* mutant phenotype in ABA-induced stomatal closure. Similar to the CSD1-GFP line, GC-CSD1-GFP seedlings grown on either 2 μM or 0.01 μM Cu closed their stomata in response to ABA (Fig. 5E-G). These findings support the conclusion that the GC CSD1 pool contributes to ABA-induced stomatal closure, regardless of SOD activity.

### Altered ROS accumulation in response to ABA in guard cells of csd1

ABA-induced stomatal closure relies on coordinated ROS production across multiple cellular compartments, and disruption of ROS generation impairs this process (da Silva et al., 2025). Accordingly, we monitored ROS accumulation in GCs after ABA treatment using fluorescence microscopy with the 2′,7′-dichlorodihydrofluorescein diacetate (H₂DCFDA) and CellROX Green fluorescent ROS probes. ABA treatment led to a substantial increase in fluorescence signal intensity of both probes in stomata of Col-0 (Fig. 6A, D, E). Instead, *csd1* mutants failed to accumulate ROS, with signal intensities showing no response to ABA treatment (Fig. 6B-E). This observation indicates that a lack of CSD1 disrupts ABA-dependent stomata closing by impairing ROS production and therefore, acts probably upstream of the ROS burst in the ABA signaling cascade.

**Figure 6.**
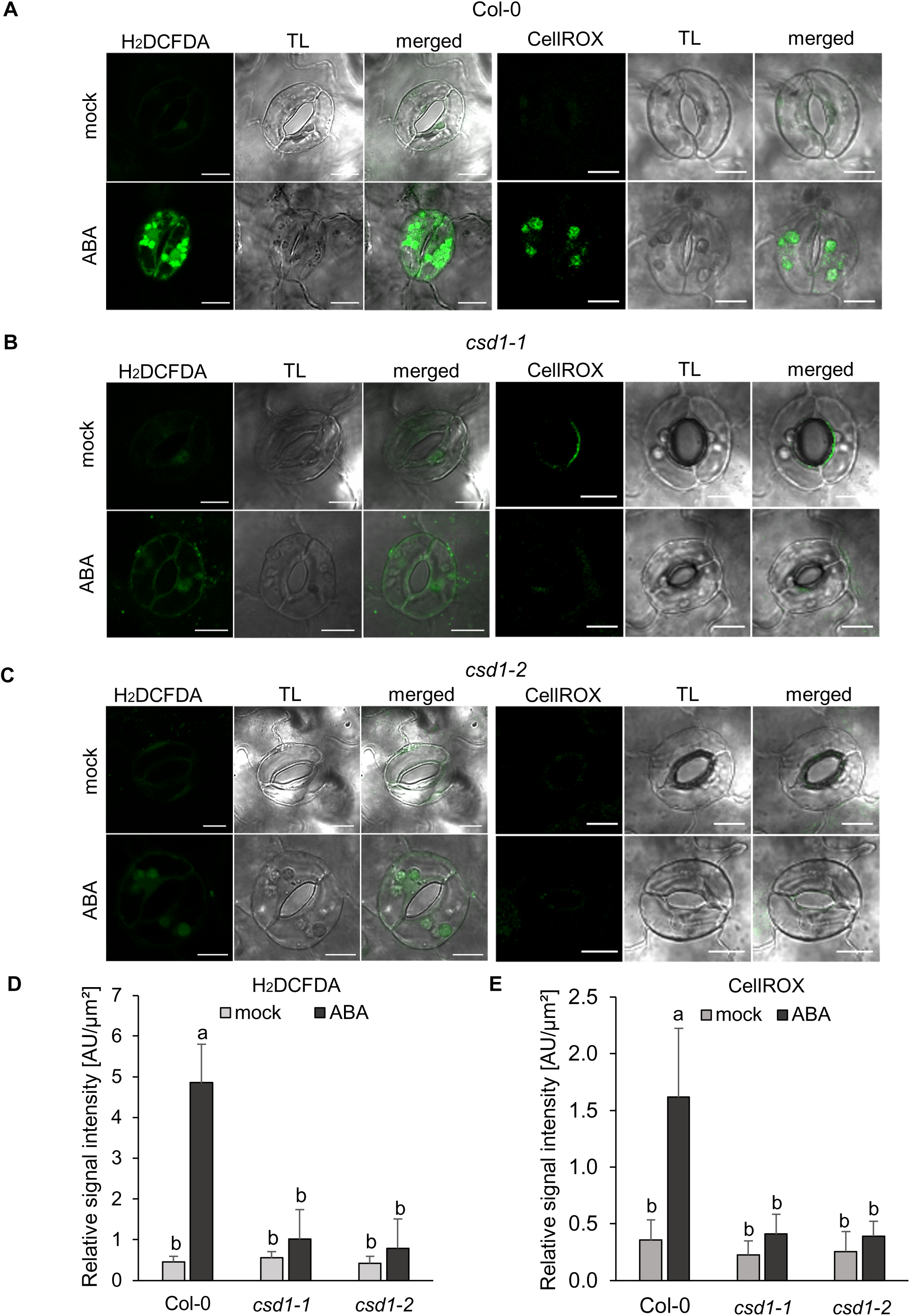
ROS distribution in guard cells of *Arabidopsis* Col-0 and *csd1* mutants. **(A-C)** Representative images showing H2DCFDA and CellROX autofluorescence localized mainly in the chloroplast in the mock and ABA-treated guard cells of (A) Col-0, (B) *csd1-1*, and (C) *csd1-2* lines grown on 2 µM Cu 1/2MS medium. TL – transmitted light. **(D-E)** Quantification of the relative signal intensity of (D) H2DCFDA and (E) CellROX probes. Signal intensity from the whole stomata (AU) was normalized to the measured stomatal area (µm²) and expressed as the mean ± SD (n = 20). Levels not connected by the same letter differ significantly (one-way ANOVA, P<0.01). Scale bar – 10 μm.

## Discussion

Soil Cu concentrations typically range from 6 to 80 mg/kg, yet bioavailability is strictly governed by soil physicochemical properties (Xu et al., 2024). Solubility is enhanced in acidic conditions but limited in alkaline, clay-rich, or high-organic-matter soils. While anaerobic or waterlogged conditions can temporarily sequester Cu as sulfide precipitates, fluctuations in pH or drainage can rapidly remobilize the metal. These environmental factors drive frequent and rapid shifts in Cu availability for plant uptake (Chia et al., 2025). Consequently, plants have to respond to them rapidly and reorganize their Cu-dependent metabolism. At the molecular level, these acclimations involve the redistribution of Cu from CSDs and CCS, as well as a functional exchange with FSD1 in *Arabidopsis*. The kinetics and extent of such changes could be especially critical under (a)biotic stress conditions, potentially determining how effectively plants cope with abrupt fluctuations in Cu availability. For this reason, a full understanding of the roles and subcellular localizations of Cu-dependent SODs remains essential.

At the cellular level, we demonstrate that CSD1 localizes not only to the cytosol but also to the nuclei of *Arabidopsis* cells. The nuclear localization of CSD1-GFP cannot be explained solely by passive diffusion, as our FRAP analysis revealed that most of the nuclear CSD1-GFP pool is immobile. Similar nuclear localization was observed for yeast *Sc*SOD1, which can relocate to the nucleus to activate the stress-response transcription factor Mec1 (Dialynaki et al., 2023); and for human *Hs*SOD1, which regulates expression of ROS-responsive genes (Tsang et al., 2014; Tsang et al., 2018; Li et al., 2019) and modulates the glucose-sensing pathway (Reddi and Culotta, 2013). Based on the high conservation of Cu/ZnSOD proteins in different organisms (Dreyer and Schippers, 2019), we propose that *Arabidopsis* CSD1 may have functions beyond its canonical antioxidant role. Moreover, because FSD1 shares the same cellular localization, its cytoplasmic and nuclear-specific roles might be complemented by CSD1, while its functions within the chloroplast or plastid may be compensated by CSD2 (Melicher et al., 2022).

Surprisingly, the dynamics of Cu-dependent functional transition between FSD1 and CSD1 have not yet been investigated. We selected 0.01 µM Cu as a limiting concentration near the threshold for permissive *Arabidopsis* growth and 2 µM Cu as a non-toxic concentration that fully supports the Cu-dependent regulation of the abundance and activity of FSD1 and CSDs. (Wang et al., 2022). Our results highlight a rapid and dynamic Cu-dependent FSD1/CSD1 switching mechanism, initiated within 20-30 h and fully restored within 2-3 days. In most cases, changes in protein abundance were accompanied by proportional changes in activity, and *vice versa*. However, under decreased Cu availability, FSD1 activity remained unchanged at 1DAT despite its reduced abundance, contradicting the general correlation. Thus, we hypothesize that under elevated Cu, FSD1 abundance decreases, while the activity of the residual protein pool may be enhanced to maintain the required level of total SOD activity until the CSD pool increases sufficiently. The mechanism of such activation is unknown and unexplored. Recently, we described the role of RECEPTOR FOR ACTIVATED PROTEIN KINASE C 1A (RACK1A) as a protein negatively modulating FSD1 activity (Melicher et al., 2026). In addition, RACK1A may negatively regulate miR398 levels, ensuring simultaneous regulation of CSDs and CCS abundance (Speth et al., 2013). All these findings eventually may suggest a role for RACK1A in regulating Cu-dependent SODs.

On the other hand, CSD1 protein abundance reacts more slowly to declining Cu level, in comparison to FSD1 response to increasing Cu availability. This difference likely reflects distinct regulatory pathways. Cu status is sensed by SPL7, which directly induces *FSD1* expression, whereas repression of *CSDs* occurs indirectly through the induction of *MIR398s* (Yamasaki et al., 2009). Another possible explanation may involve the buffering capacity of cellular Cu stores (Burkhead et al., 2009), which could temporarily maintain CSDs and CCS protein levels as Cu availability declines. Additionally, it was shown that the *SPL7* transcript level remains relatively constant regardless of Cu availability (Bernal et al., 2012). Thus, SPL7 is thought to sense Cu through post-translational mechanisms (Garcia-Molina et al., 2014). Our results are consistent with this, as the response of SPL7-dependent proteins occurs within hours. Therefore, residual Cu availability may inhibit full activation of SPL7, as well as expression of *FSD1* and the regulatory *MIR398s*.

Despite the tight regulation, the stomatal pool of CSD1 did not respond to changes in Cu content. This pattern was not observed in FSD1, highlighting a specific form of regulation unique to CSD1. Cu-dependent regulation of CSD1 occurs through the miR398-mediated downregulation. Its BS at the *CSD1* sequence is located in the 5′UTR, where two parts are disrupted by an intron (Fig. 4B; Fig. S12A). Consistently, under drought stress, peanut *AhCSD1* has been reported to produce an alternatively spliced form lacking the miR398 BS (Park and Grabau, 2017). This event is likely conserved across all plant isoforms homologous to CSD1, as such splicing variants have been detected in *Glycine max* (Park and Grabau, 2017), *Solanum lycopersicum*, *Oryza sativa*, and *Zea mays* (Fig. S15A-C). The presence of an intron that disrupts the miR398 binding site was also identified in *Medicago truncatula* and *Solanum tuberosum* (Fig. S15D), though spliced variants have not yet been identified. Despite this evidence, the function of the miR398-insensitive *CSD1.2* splice variant across plant species requires further research. Aside from this, miR398 targets another three mRNAs: *CSD2*, *CCS*, and a *SUBUNIT OF THE MITOCHONDRIAL CYTOCHROME C OXIDASE* (*COX5b-1*) in *Arabidopsis* (Jones-Rhoades and Bartel, 2004; Bonnet et al., 2004; Sunkar et al., 2006; Beauclair et al., 2010). Interestingly, the miR398 BS of *CCS* and *CSD2* is located within their coding sequence (Bonnet et al., 2004; Beauclair et al., 2010). This region is conserved among all CSDs and CCS on the amino acid level in *Arabidopsis* (Fig. S16A) and presumably also in other crop species. However, the nucleotide sequence of *CSD1* differs just enough to limit miR398 binding (Fig. S16B). These observations suggested that selective pressure favored synonymous substitutions in CSD1 that weakened miR398 binding without altering the encoded protein sequence, whereas the presence of BS in the 5′UTR allowed the selective bypass of miR398 regulation.

So far, no direct evidence demonstrates the involvement of SODs in ABA-mediated stomatal closure. Their contribution has been inferred from their role in ROS metabolism, particularly in catalyzing the conversion of O_2_^•–^ to H_2_O_2_ produced by RBOHs or other O_2_^•–^ sources (Sierla et al., 2016; Rodrigues and Shan, 2022). Here we show that both *csd1* mutant lines exhibited impaired ABA-induced stomatal closure with direct physiological impact. In contrast, all complemented lines (CSD1-GFP, CSD1m-GFP, GC-CSD1-GFP) restored the phenotype regardless of Cu. Concurrently, under low-Cu conditions, none of these lines showed detectable CSD1 enzymatic activity. This is consistent with the absence of CCS, a key chaperone required for Cu^1+^ insertion into the active site and for proper disulfide bond formation (Brown et al., 2004; Furukawa et al., 2004). Nonetheless, studies of *Hs*SOD1 have shown that, even in the absence of Cu insertion, the SOD1 polypeptide can adopt a partially mature, monomeric form (Carroll et al., 2004), which may retain non-catalytic functions. Although the proof of the functionality of this form in *Arabidopsis* requires further in-depth study, we suggest that CSD1’s role in stomatal closure depends more on the protein itself than on its enzymatic activity. The observation that ABA treatment did not trigger substantial ROS production in the *csd1* mutant, as it did in Col-0, suggests that the ABA-induced stomatal closure in the mutants is disrupted upstream of ABA-induced ROS production. Hence, compartmentalized ROS production is considered a crucial step in proper stomatal closure (da Silva et al., 2025). However, the exact role of CSD1 in this pathway remains unclear.

In conclusion, CSD1 and FSD1 alternate in a Cu-dependent manner. However, their distinct subcellular, tissue, and organ localizations, as well as the dynamics of their Cu-dependent exchange, suggest isoform-specific roles. CSD1 plays a unique, enzyme-independent role in stomatal closure, driven by its guard cell-specific localization which occurs because a cell type-specific splice variant (*CSD1.2*) carries an altered binding site that escapes miR398-mediated regulation (Fig. 7). Our findings expand the functional range of CSD1 beyond traditional antioxidant activity, opening new avenues for investigating its broader physiological role in stomatal regulation during essential plant development and stress responses.

**Figure 7.**
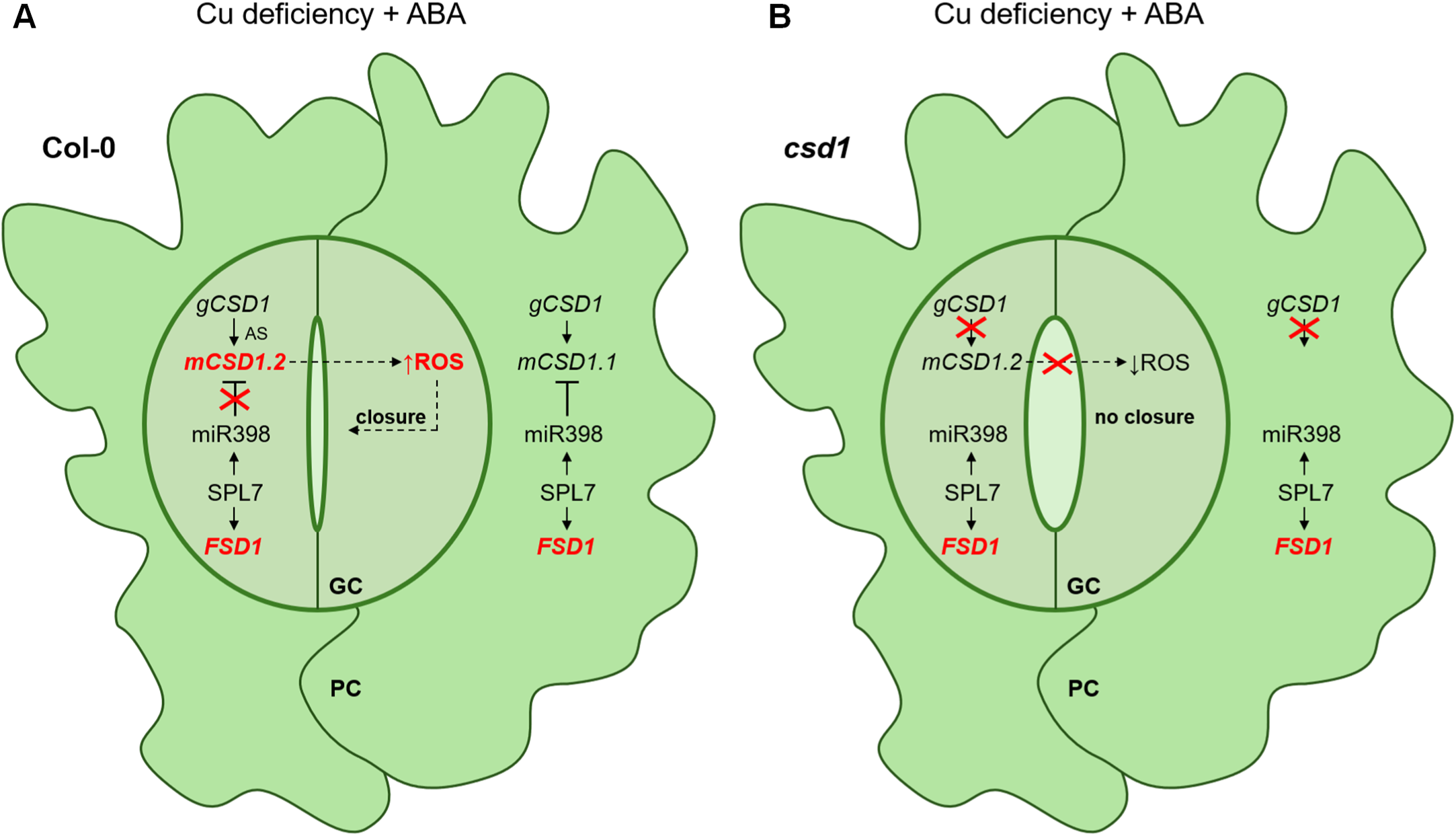
Proposed model of *CSD1* alternative splicing in guard cells during ABA-mediated stomatal closure. **(A)** Under copper-deficient conditions in Col-0, the transcription factor SQUAMOSA PROMOTER-BINDING-LIKE PROTEIN 7 (SPL7) induces the expression of *IRON SUPEROXIDE DISMUTASE 1* (*FSD1*) and the regulatory *MIR398* in both pavement cells (PCs) and guard cells (GCs). The *COPPER/ZINC SUPEROXIDE DISMUTASE 1* gene (*gCSD1*) generates the normally spliced transcript (*mCSD1.1*) in PCs, where it is downregulated by mature miR398. In contrast, alternative splicing (AS) in GCs produces the transcript variant *mCSD1.2*, which contains an altered miR398-binding site and is therefore insensitive to miR398-mediated regulation. Upon elevation of abscisic acid (ABA) levels, translated CSD1.2 indirectly contributes to the generation of a reactive oxygen species (ROS) burst, promoting stomatal closure. **(B)** In the *csd1* mutant, the miR398-insensitive m*CSD1.2* variant is absent, disrupting ABA signaling and preventing the ROS burst. Consequently, stomata fail to close in response to ABA.

## Methods

### Plant materials and growth conditions

*Arabidopsis thaliana* wild-type plants, ecotype Columbia (Col-0), *fsd1-1* (Dvořák et al., 2021), *csd1-1* (SALK_109389), and *csd1-2* (SALK_024857), both obtained from the European Arabidopsis Stock Centre (http://arabidopsis.info/BasicForm), and a previously prepared transgenic line carrying *pFSD1::GFP:FSD1-3′UTR-FSD1* (FSD1-GFP) (Dvořák et al., 2021) and a *35S::sGFP* construct for free GFP cloned using pMAT037 plasmid (Matsuoka and Nakamura, 1991; Mano et al., 1999) were used in this study. The *fsd1-1/csd1-1 (fsd1/csd1)* double mutants were generated by genetic crossing and selected using PCR genotyping and immunoblotting. All primers used to verify mutants are listed in Supplementary Table 1.

Surface-sterilized *Arabidopsis* seeds were stratified at 4 °C for 1-2 days to synchronize germination and then were grown on homemade 1/2 Murashige and Skoog (MS) medium, adjusted to final CuSO_4_·5H_2_O concentrations of 0.01 μM and 2 μM (hereafter referred to as 0.01 μM Cu and 2 μM Cu, respectively), and solidified with 0.5% (w/v) Phytagel. Seedlings were grown vertically at 21°C, 16/8 h (light/dark) photoperiod with an illumination intensity of 150 μmol photons‧m^−2^‧s^−1^ in a phytochamber (Weiss Technik) for 1-15 days, depending on the purpose. Additionally, the above-mentioned lines and newly prepared transgenic lines were grown from seeds in soil-filled pots for 4 weeks, or until final propagation, under the same conditions as described above.

### Preparation of constructs and transgenic lines

The C-terminal fusion construct of *eGFP* (hereafter designated as GFP) with genomic DNA (gDNA) of *CSD1* under its native promoter (1731 bp upstream of the start codon) *pCSD1::CSD1:GFP::3′UTR-CSD1*, was cloned using the MultiSite Gateway Three-Fragment Vector Construction kit (Thermo Fisher Scientific, USA) as a cloning method. For the 3′UTR, 511 bp downstream of the stop codon was used. The amplified promoter and the genomic DNA sequence were recombined into the pDONRP4-P1R vector (fragment A) and the 3′UTR into the pDONRP2R-P3 donor vector (fragment C) (Supplementary Table 1). The plasmid pEN-L1-F-L2, which contains a *GFP* with a stop codon, served as fragment B for the subsequent three-fragment LR recombination into the destination vector pB7m34GW0.

The C-terminal fusion construct of GFP with gDNA of *CSD1* under a GC-specific promoter (*pGC1::CSD1:GFP::3′UTR-CSD1*) was cloned using Gibson assembly. The fusion construct CSD1-GFP was used as a template for a PCR reaction with primers containing overhangs for the GC-specific promoter pGC1 (Gibberellin-regulated protein 9; At1g22690) (Supplementary Table 1).

The C-terminal fusion construct of GFP with gDNA of *CSD1* under a native promoter with five point mutations in the miR398 binding site (*pCSD1m::CSD1:GFP::3′UTR-CSD1*) was cloned using Gibson assembly. The nucleotides that were mutated were chosen according to the previously published study (Dugas and Bartel, 2008). The CSD1-GFP fusion construct was used as a template for a PCR reaction with primers containing a linker and an added mutation in the miR398 binding site (Supplementary Table 1).

Constructs of *GFP* reporter with nuclear localization signal (*^NLS^GFP*) driven by the native promoters of *MIR398a*, *MIR398b*, and *MIR398c* were prepared using Gibson assembly. Consequently, promoter sequences of all *MIR398*s (3000 bp upstream of the start codon) were inserted into the Gateway Dual Selection Vector pENTR2B-NLS-GFP (Hajný et al., 2024), using primers with overhangs (Supplementary Table 1). A one-component LR reaction was used to insert the cassette into the pMCS-GW destination vector.

The cloned products, validated by sequencing, were transformed into *Agrobacterium tumefaciens* GW3101 for further stable transformation of the *csd1-1* and *csd1-2* mutants, generating transgenic lines carrying *pCSD1::CSD1:GFP::3′UTR-CSD1* (hereafter designated as CSD1-GFP); *pCSD1m::CSD1:GFP::3′UTR-CSD1* (hereafter designated as CSD1m-GFP), and *pGC1::CSD1:GFP::3′UTR-CSD1* (hereafter designated as GC-CSD1-GFP). *Agrobacterium* bearing constructs of *pMIR398::^NLS^GFP* were used for stable transformation of Col-0, generating *pMIR398a::^NLS^GFP*, *pMIR398b::^NLS^GFP*, and *pMIR398c::^NLS^GFP* transgenic lines.

### Protein extraction, immunoblotting and in-gel SOD activity analysis

To prepare the protein extract, the plant material homogenate was resuspended in the native extraction buffer (50 mM sodium phosphate (pH 7.8), 10% (v/v) glycerol, 2 mM ascorbate), placed on ice for 30 min, and vortexed occasionally. The extract was centrifuged at 15,000×g for 20 min at 4 °C, and the protein concentration of the supernatant was measured. The protein extract was used for the in-gel SOD activity assay and total SOD activity. For immunoblot analysis, native protein extracts were supplemented with a 3:1 ratio of 4× concentrated Laemmli SDS buffer to 5% (v/v) β-mercaptoethanol. The samples were boiled for 5 min at 95 °C before SDS-PAGE electrophoresis.

The immunoblotting and in-gel activity analyses were performed according to the previously used protocol (Melicher et al., 2022). Primary antibodies such as anti-CSD1/2 (Agrisera AS06170; dilution 1:2000 in TBS-T with 1% (w/v) low-fat dry milk), anti-FSD1 (Agrisera AS06125; dilution 1:2000 in TBS-T with 1% (w/v) low-fat dry milk), and CCS (Agrisera AS07219; dilution 1:2000 in TBS-T with 1% (w/v) low-fat dry milk) were used for specific SOD inhibition assays, gels were preincubated for 15 min in 50 mM sodium phosphate buffer (pH 7.8) containing either 5 mM H_2_O_2_ (to inhibit CSDs and FSD1) or 2 mM KCN (to inhibit only CSDs) in 50 mM sodium phosphate buffer (pH 7.8).

### Experimental setup for SODs and CCS Cu-response dynamics

CSD1-GFP and FSD1-GFP lines were cultured under normal growth conditions on 0.01 μM Cu or 2 μM Cu for 3 days. One-third of them were selected for further analysis (sample “before”), and the other two-thirds were transferred to a medium with the same Cu concentration (batch “control”) or with the opposite concentration (batch “transfer”). Each seedling was imaged using an Axio Zoom.V16 stereomicroscope (Carl Zeiss, Germany) before transfer and on the 1^st^, 2^nd^, and 3^rd^ day after transfer (DAT), using the same microscope settings. GFP signal intensity was measured from micrographs using Zeiss ZEN software (Black and Blue versions, Carl Zeiss). Regions of interest (ROI) were drawn around individual hypocotyls, and the integrated fluorescence density in arbitrary units (AU) was extracted. To account for size differences among ROIs, the total GFP signal was normalized to ROI area (µm^2^), yielding area-normalized fluorescence intensity for all comparisons. The experiment was performed in triplicate for each line and condition (n = 50) to evaluate statistical significance using a one-way ANOVA with post hoc Tukey HSD test. Representative images of the cotyledons were taken separately using a confocal laser scanning microscope LSM710 (Carl Zeiss, Germany).

In parallel, Col-0 seedlings (n = 90) were used under the same experimental conditions to determine the abundance of CSD1, CSD2, FSD1, and CCS by immunoblotting and in-gel SOD activity, as described above.

### Measurements of stomatal aperture, leaf stomatal conductance (gsw), and leaf temperature

7-day-old seedlings were incubated in a stomatal open buffer (50 mM KCl, 0.1 mM CaCl_2_, and 10 mM MES, pH 6.15) under the light (150 μmol photons‧m^−2^‧s^−1^) for 2.5 h, then transferred to the stomatal open buffer with or without 50 μM ABA for 3 h. Afterward, seedlings were placed on a glass slide in a drop of open buffer and covered with a coverslip. Stomata were imaged with an Axio Imager Z2 microscope (Carl Zeiss, Germany) using an EC Plan-Neofluar ×40/0.75 M27 objective in bright-field conditions. Stomatal aperture was measured as a ratio of aperture width and length, which is reduced upon closure, in ImageJ (Broken Symmetry Software) across three independent replicates (n = 50). Statistical significance was evaluated using a one-way ANOVA with post hoc Tukey HSD test.

4-week-old plants of Col-0, *csd1-1*, *csd1-2*, and CSD1-GFP were cultivated in soil under controlled conditions as described above. Plants were sprayed 2 h after the onset of the light phase with either distilled water containing 0.01% (v/v) Tween-20 (mock treatment) or the same solution supplemented with 100 μM ABA. Following application, plants were maintained in the phytochamber under the same growing light conditions for an additional 1 h. Leaf gsw was measured using the LI-600 porometer/fluorometer (LI-COR Environmental, Lincoln, NA, USA) while the plants were still under growing conditions. For each genotype, the gsw was estimated from 50-80 leaves from at least 20 individual plants (n = 20). Statistical significance was evaluated using one-way ANOVA with post hoc Tukey HSD test. Afterward, leaf temperature was estimated from thermal images of plants captured with a thermal camera, Optris Xi 400 (field of view 80°×54°) (Optris, Berlin, Germany), using Process Imager Software, Optris PIX Connect. The thermal camera operates in the long-wave infrared spectrum (8-14 µm) with a 382×288-pixel resolution. For each variant, three plants were scanned, and 20 temperature values (averages from a 5×5-pixel area) were assessed per plant (n = 60). Statistical significance was evaluated using one-way ANOVA with post hoc Tukey HSD test.

### RNA isolation and RT-PCR analysis of splicing

RNA was isolated from 14-day-old leaves of Col-0 grown on 0.01 μM or 2 μM Cu. RNA isolation and cDNA preparation were performed according to the previously used protocol (Hrbáčková et al., 2021). The obtained cDNA was used directly for RT-PCR analysis of splicing events in CSD1. Each 50 μl PCR reaction contained two forward primers and one reverse primer, yielding products of 512 bp and 402 bp, respectively. Detection of splice and full variants was performed in 3 biological replicates, with 10 seedlings per replicate (n = 30). Primer sequences are listed in Supplementary Table 1.

### Enrichment of guard cell fraction and measurement of total SOD activity

The GC-enriched fraction was obtained according to the previously used protocol (Brosche, 2017). Briefly, leaves from 14-day-old GC-CSD1-GFP and *csd1-1* mutant seedlings grown on 0.01 μM Cu or 2 μM Cu were cut and blended with 250 μl of Milli-Q cold water and crushed ice for 1 min. The obtained amount was passed through a nylon mesh (100 μm pore size), and the pellet was collected for subsequent homogenization in liquid nitrogen. Together with homogenized 14-day-old leaves of GC-CSD1-GFP grown on 0.01 μM or 2 μM Cu, protein extracts were obtained and designated as total (T) and enriched (E), and were used for immunoblotting and in-gel visualization of SOD activity. The total SOD activity was measured using a colorimetric method with the SOD activity assay kit (Merck, CS0009) according to the manufacturer’s protocol. Each replicate (total volume 200 μl) consisted of 20 μl of GC-enriched fraction containing 3 μg of total protein. The measurements were performed in 3 biological replicates.

### ROS staining

DCF imaging and quantification were performed according to the previously used protocol (Kim and Xue, 2020), with minor modifications. 7-day-old seedlings of *Arabidopsis* Col-0, *csd1-1*, and *csd1-2*, grown on 1/2 MS medium supplemented with 2 µM Cu, were incubated in stomatal opening buffer for 2 h under light conditions (150 μmol photons‧m^−2^‧s^−1^). Subsequently, seedlings were transferred to a fresh opening buffer containing 5 µM H₂DCFDA (Thermo Fisher Scientific, USA), either with or without 50 µM ABA, and incubated for 30 min in the dark. Following staining, GCs were imaged using a confocal laser scanning microscope (LSM710; Carl Zeiss, Germany). CellROX Green (Thermo Fisher Scientific, USA) staining was performed using the same protocol. Signal intensity was measured using Zeiss ZEN software (Black and Blue versions, Carl Zeiss). ROI were drawn around stomata, and the integrated fluorescence density was extracted in AU. To account for size differences among ROIs, the intensity in AU was normalized by the ROI area (µm²), yielding an area-normalized fluorescence intensity used for all comparisons. For each line and treatment, 10 plants were imaged (n = 20). Statistical significance was evaluated using one-way ANOVA with post hoc Tukey HSD test.

### Confocal laser scanning microscopy

5-day-old CSD1-GFP seedlings (in the *csd1-1* background) grown on 2 μM Cu were used for microscopy. Imaging was performed using a confocal laser scanning microscope LSM880 equipped with an Airyscan detector (Carl Zeiss, Germany) or LSM990 equipped with an Airyscan 2 detector (Carl Zeiss). Image acquisition was performed using 20× (0.8 NA) and 40× (1.4 NA) Plan-Apochromat objectives. Samples were imaged with a 488 nm excitation laser line using emission filters BP420-480 + BP495-550 for GFP detection, with the laser intensity kept below 2% of the available range. Images were processed as single-plane maximum-intensity projections of Z-stacks in Zeiss ZEN 3.13 (Blue version), assembled and finalized in Microsoft PowerPoint to produce the final figures.

### Light-sheet fluorescence microscopy

For LSFM imaging, a 4-day-old seedlings growing on either 0.01 µM or 2 µM Cu were inserted into a fluorinated ethylene propylene (FEP) tube (Wolf-Technik, Germany), with cotyledons being exposed to air, hypocotyl being embedded in 1% (w/v) agarose inside the FEP tube, and root being exposed to liquid medium inside the LSFM observation chamber (Ovečka et al., 2015). The FEP tube with the mounted seedling was attached to the holder and placed in a pre-tempered (21°C) LSFM observation chamber filled with a filter-sterilized liquid 1/2 MS medium (pH 5.8) supplemented with either 0.01 µM or 2 µM Cu. Circulation of culture medium between the reservoir of fresh medium and the LSFM observation chamber was secured by the Minipulse Evolution peristaltic pump (Gilson Inc.). The adjustable multicolor LED illumination reflector secured the 16 h light/8 h dark cycle during long-term live-cell imaging. After sample insertion and stabilization, long-term developmental live-cell imaging was performed on a Zeiss Lightsheet Z.1 (Carl Zeiss, Germany) with dual-side illumination using two 10×/0.2 NA illumination objectives (Carl Zeiss, Germany) and a pivot scan mode, with an excitation laser line at 488 nm, a beam splitter LP560, and an emission filter BP 505-545. Image acquisition was performed with a W Plant-Apochromat 10×/0.5 NA water-immersion detection objective (Carl Zeiss, Germany) and recorded with a sCMOS camera (PCO.Edge, PCO AG) at an exposure time of 50 ms per optical section. Time-course imaging was carried out at 30 min intervals in Z-stack mode for 54−72 h. Post-processing was performed using Zeiss ZEN 3.13 (Blue version) or ImageJ.

### Fluorescence recovery after photobleaching (FRAP)

FRAP analysis in nuclei of hypocotyl epidermal cells was conducted on CSD1-GFP and GFP *Arabidopsis* transgenic lines (the latter expressing *35S::sGFP* as free GFP), as described before (Dvořák et al., 2021). Data from a total of 45 regions of interest (for free GFP) and 53 regions (for CSD1-GFP) were exported to Microsoft Excel software and normalized to absolute fluorescence intensities, where the highest intensity (prior to bleaching) was set to 1, and the lowest intensity (immediately following bleaching) was set to 0. The FRAP measurement was conducted across three biological replicates (6 seedlings per replicate).

## Author contributions

M.T., P.D., K.H., Z.K., M.Š., J.Ř. and J.S. performed experiments and analyses. M.T. and P.D. also assisted with data assessment. J.Š. provided the infrastructure. M.T. and P.D. drafted the manuscript, which was revised and edited by T.T., M.O. and J.Š., while P.D., J.Š. and M.O. secured the funding. P.D. and T.T conceived and supervised this study. All authors approved the final version of the manuscript.

