## Supplementary figures 1-16 for "Stomatal movement in *Arabidopsis* is driven by guard cell-localized and copper-insensitive CSD1 splice variant"

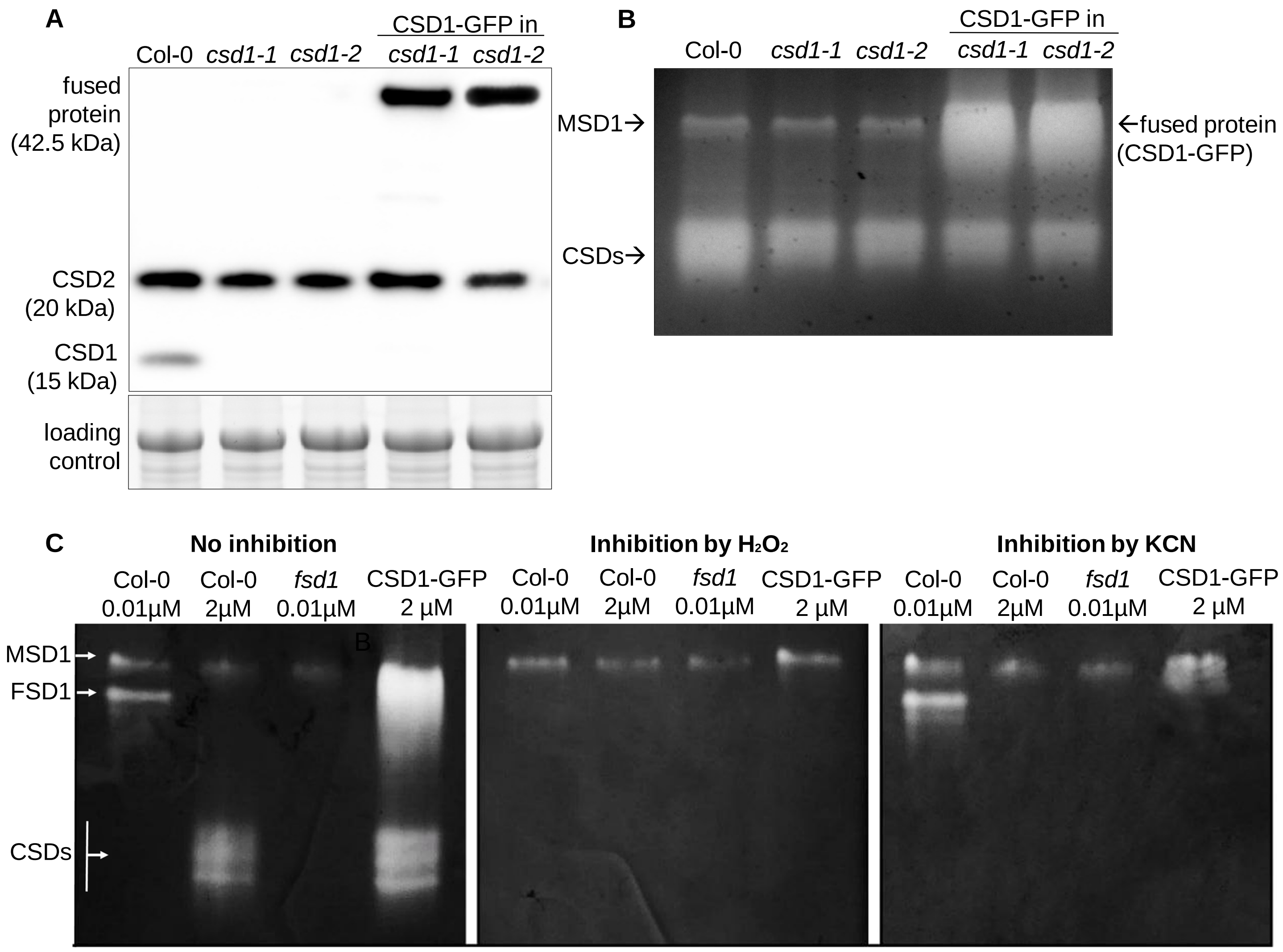


**Supplementary Figure S1. Validation of stable complemented CSD1-GFP in the background of *csd1-1* and *csd1-2* mutants. (A)** Immunoblotting analysis of CSD1 and CSD1-GFP abundance in 14-day-old *csd1* mutants, Col-0, and complemented *csd1* mutants using anti-CSD1 antibody. **(B)** Visualization of SOD isozymes on native polyacrylamide gel. **(C)** SOD inhibition assay in Col-0, *fsd1-1*, and complemented *csd1* mutants grown under different Cu availability conditions to determine the specificity of SOD isoenzymes. Native gels were analyzed without preincubation or after preincubation with KCN, which inhibits CSD activity, or $H_2O_2$, which inhibits both FSD1 and CSD activities.

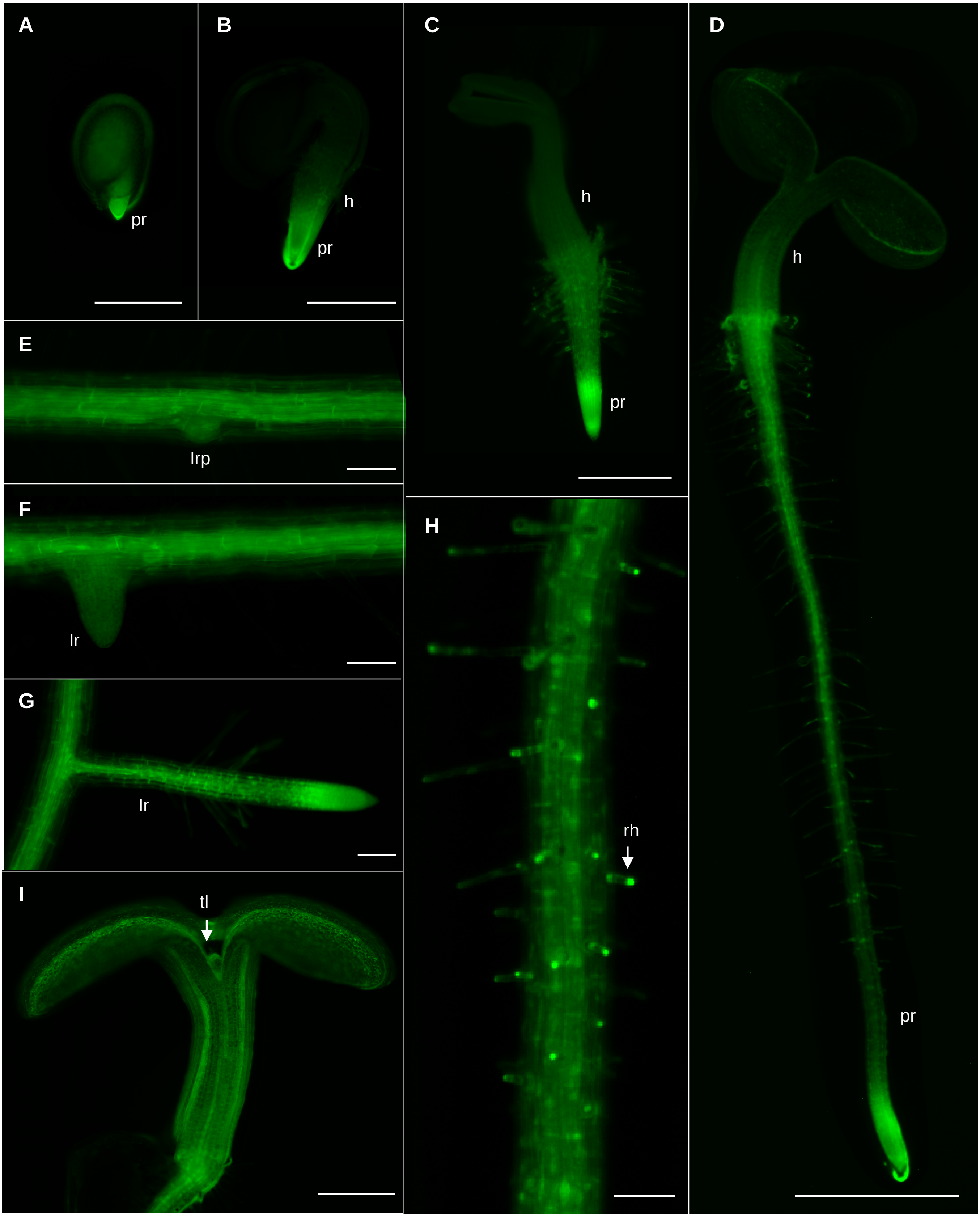


**Supplementary Figure S2. Localization of CSD1-GFP visualized by ZOOM stereomicroscope at different developmental stages of young *Arabidopsis* seedlings. (A-D)** CSD1-GFP seedling on the (A) 1$^{st}$ day after germination (DAG), (B) 2$^{nd}$ DAG, (C) 3$^{rd}$ DAG, and (D) 5$^{th}$ DAG with hypocotyl (h), primary root (pr). **(E-G)** Detailed images of lateral root development with (E) lateral root primordia (lrp), (F) developing and (G) developed lateral root (lr) on 7$^{th}$ DAG. **(H)** 8-day-old primary root with root hairs (rh). **(I)** Closer image of the aerial part of a 5-day-old seedling with fully opened cotyledons and emerging true leaves (tl). Scale bar (A-C, E) 500 µm, (D) 1000 µm (F, G-I) 100 µm.

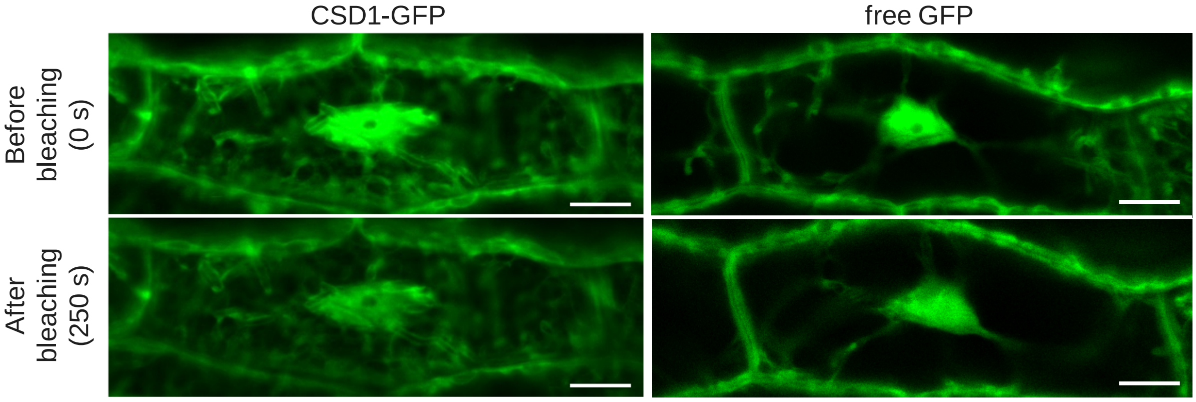


**Supplementary Figure S3.** Representative images of CSD1-GFP and free GFP nuclei before and after 250 s of selective bleaching. Scale bar – 5 μm.

A

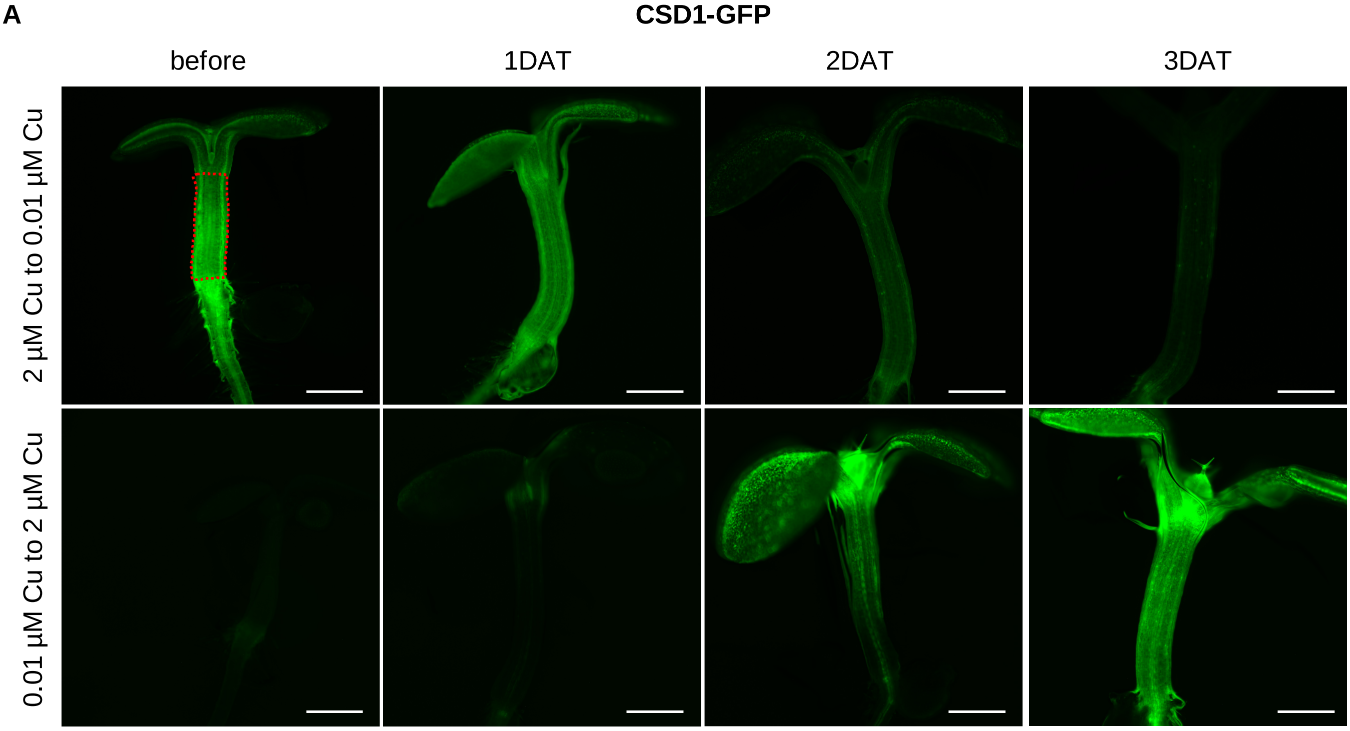


B

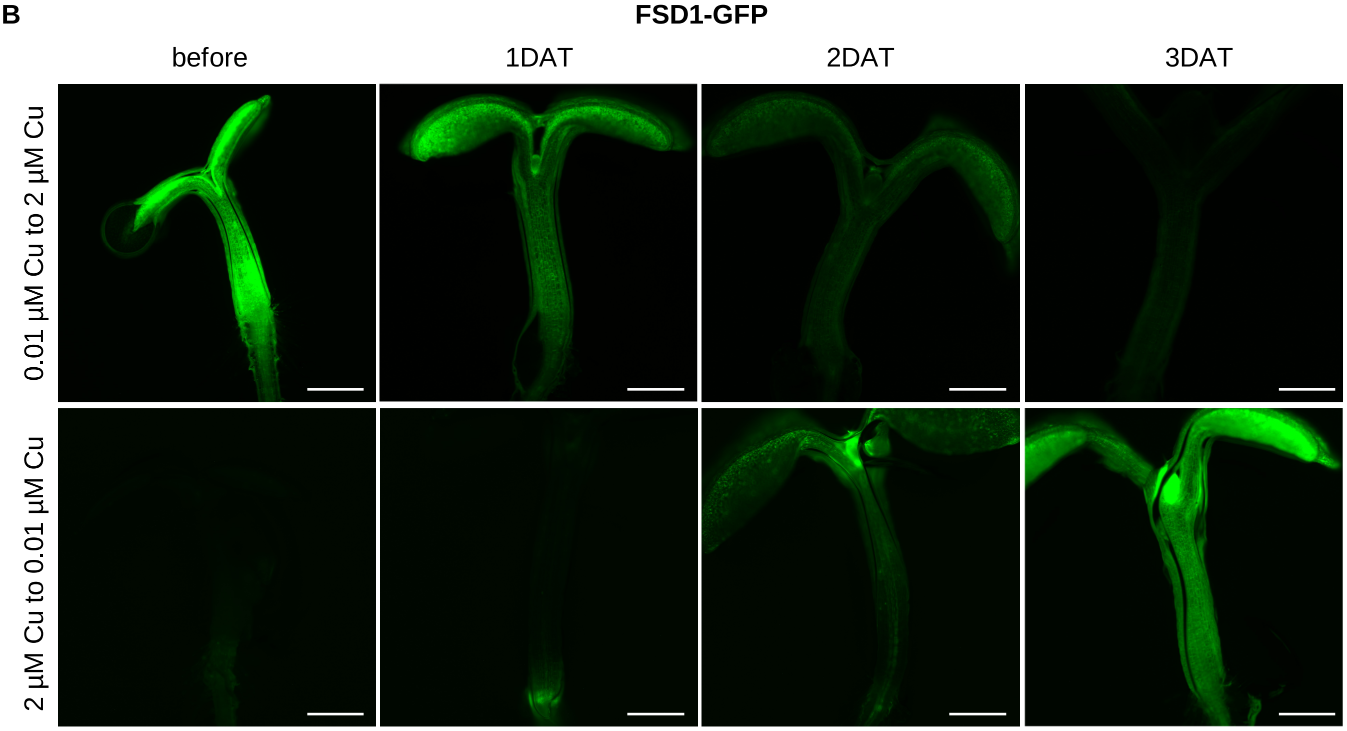


**Supplementary Figure S4. Response of complemented CSD1-GFP and FSD1-GFP lines to changes in Cu concentration.** Representative images from an Axio Zoom.V16 stereomicroscope of the **(A)** CSD1-GFP line and **(B)** FSD1-GFP seedlings before and during three days after transferring (DAT) from 0.01 µM to 2 µM Cu and from 2 µM to 0.01 µM Cu, which were used for further quantification of GFP signal intensity. ROIs were drawn around the hypocotyls, as shown by the dashed red line. Scale bar – 500 µm.

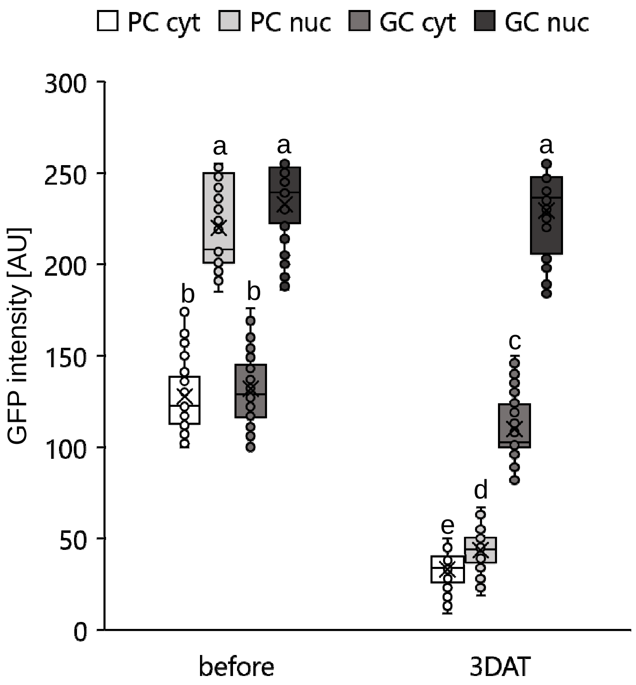


**Supplementary Figure S5. Difference in the CSD1-GFP signal intensity between the cytoplasm and nuclei of leaf pavement and stomata guard cells in response to Cu changes.** Changes in fluorescence intensity in the cytoplasm (cyt) and nuclei (nuc) of pavement cells (PC) and guard cells (GC) of the CSD1-GFP line before (2 μM Cu) and after 3DAT (0.01 μM Cu). Values are extracted in arbitrary units (AU). The experiment was performed in triplicate, with a total of n = 20 measured values. Levels not connected by the same letter differ significantly (one-way ANOVA, $P<0.01$).

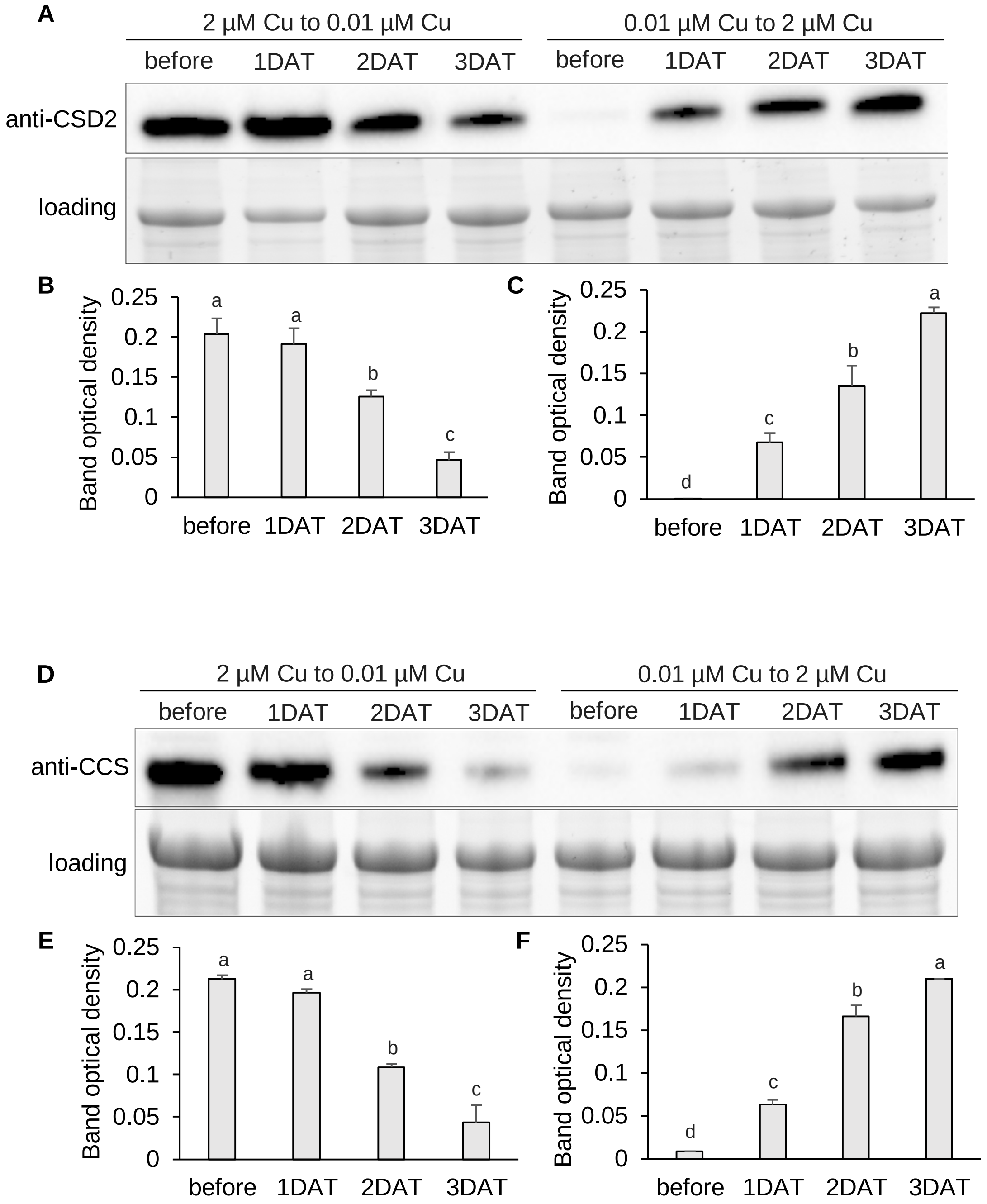


**Supplementary Figure S6. Cu-dependent changes in endogenous CSD2 and CCS abundance in *Arabidopsis*.** Visualization and statistical evaluation of **(A-C)** CSD2 and **(D-F)** CCS abundance before and during three days after transferring (1DAT, 2DAT, 3DAT) seedlings from 2 µM to 0.01 µM Cu (B, E) and from 0.01 µM to 2 µM Cu (C, F). The experiment was performed in triplicate, with a total of n = 90 examined seedlings. Error bars represent standard deviation. Levels not connected by the same letter differ significantly (one-way ANOVA, $P<0.01$).

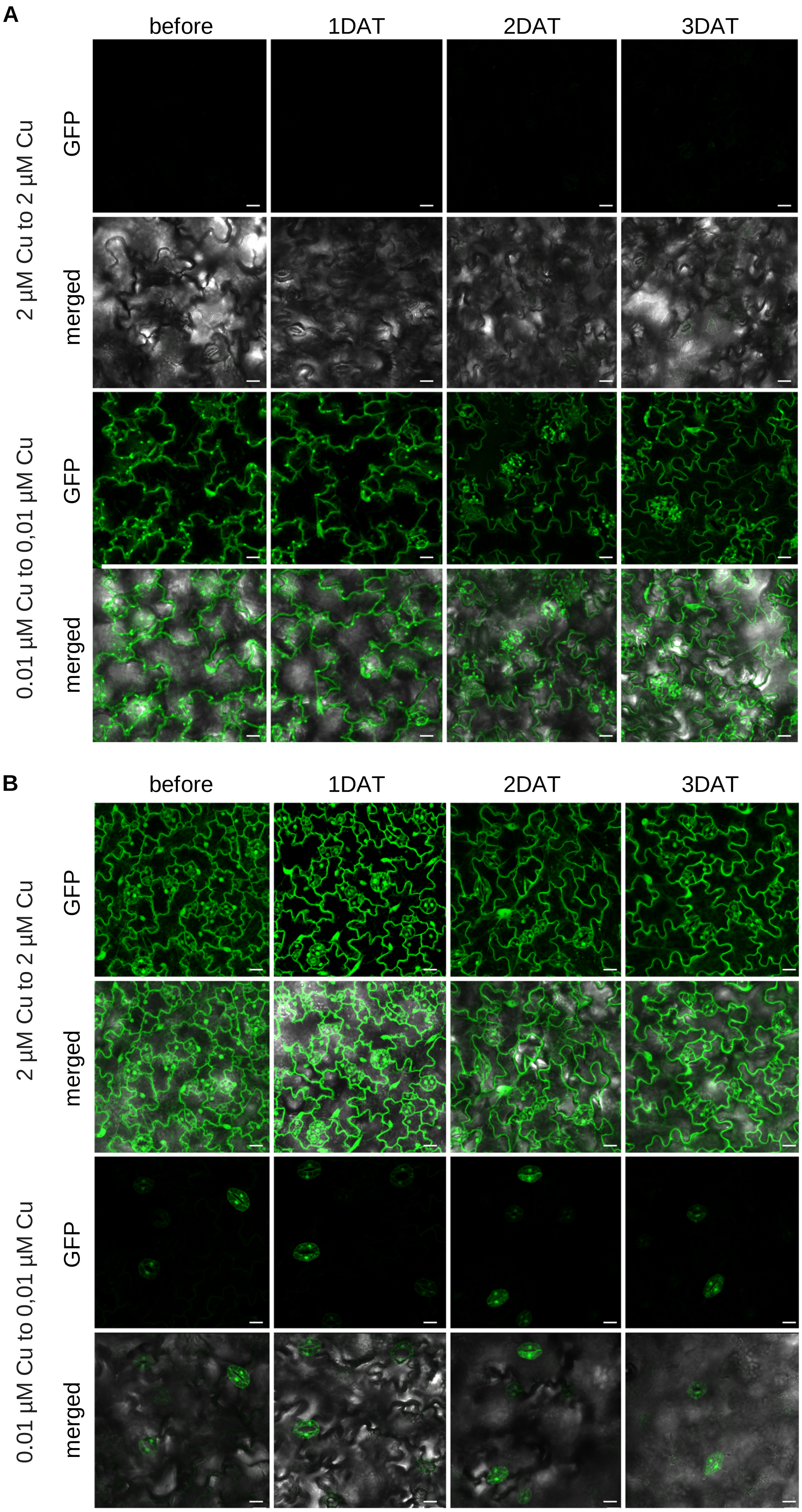


**Supplementary Figure S7. Response of complemented FSD1-GFP and CSD1-GFP lines to unchanged Cu concentration. (A-B)** Representative images of the cotyledons of (A) the FSD1-GFP line and (B) the CSD1-GFP line before and during three days after transferring (DAT) seedlings from 2 µM to 2 µM Cu and from 0.01 µM to 0.01 µM Cu. Scale bar – 20 µm.

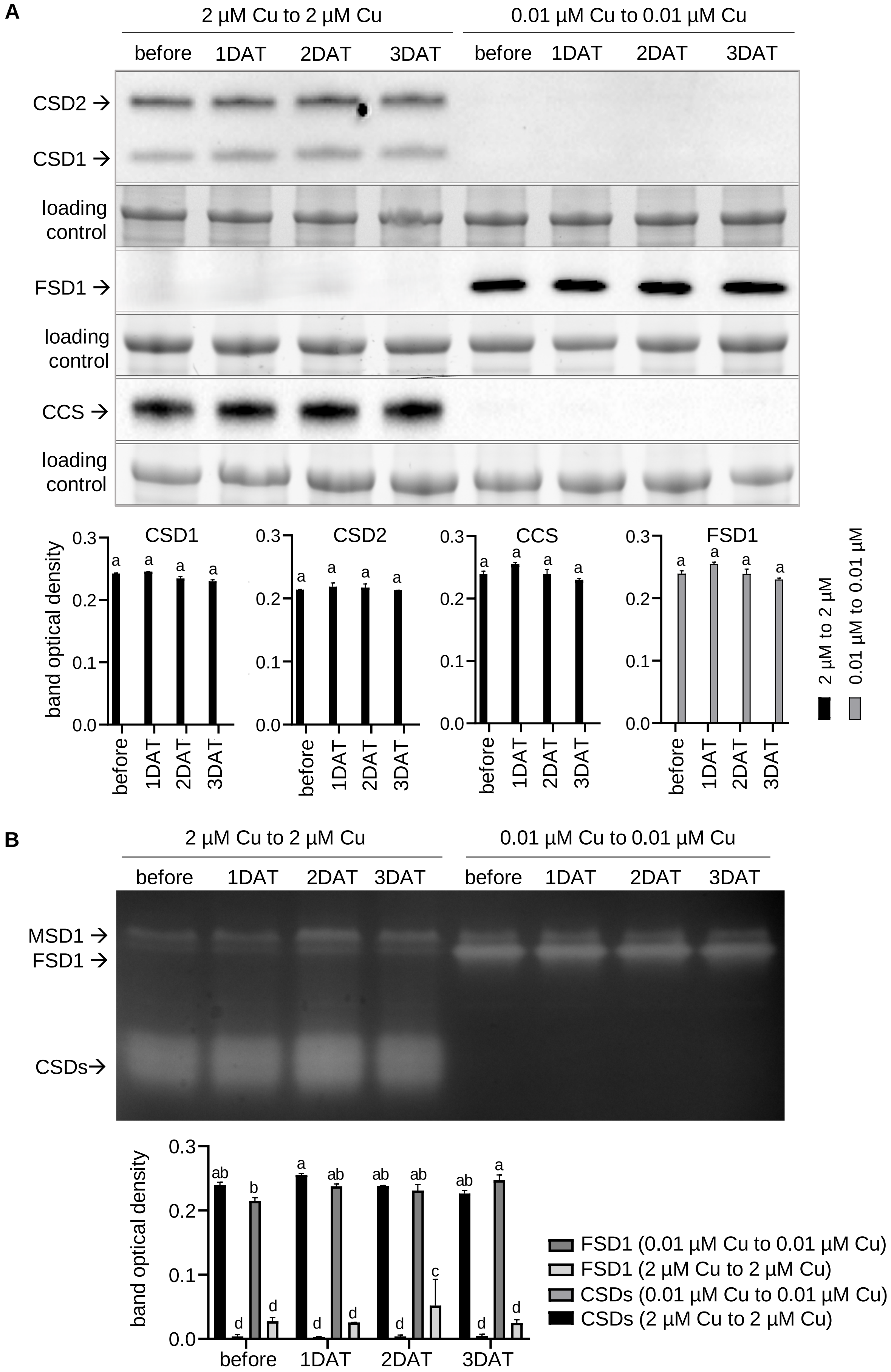


**Supplementary Figure S8. Endogenous SOD abundance and activity in control transfers in *Arabidopsis*. (A)** Abundance of CSD1, CSD2, FSD1, and CCS before and during three days after transferring (DAT) seedlings from 2 µM to 2 µM Cu and from 0.01 µM to 0.01 µM Cu. **(B)** Visualization of SOD isozymes on native polyacrylamide gel before and during 1-3DAT control seedlings. The experiment was performed in triplicate, with a total of n = 90 examined seedlings. Levels not connected by the same letter differ significantly (one-way ANOVA, P<0.01).

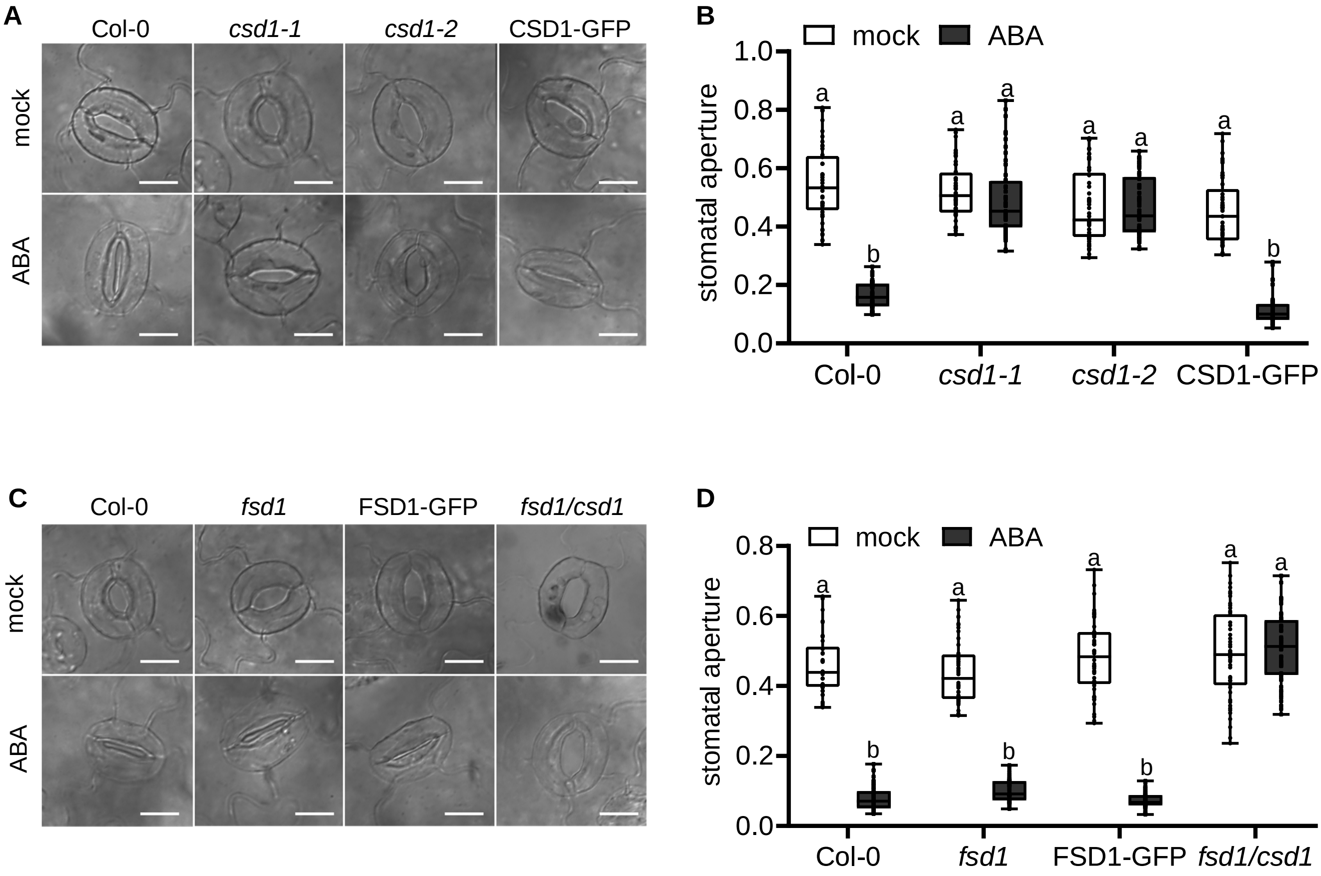


**Supplementary Figure S9. The *csd1* mutant displays dysregulated ABA-induced stomatal closure independent of copper availability.** Stomatal aperture of Col-0, single *csd1* and *fsd1*, and double *fsd1/csd1* mutants and complemented CSD1-GFP and FSD1-GFP lines was measured after application of 50 µM abscisic acid (ABA). **(A, C)** Representative images of stomata in 7-day-old seedlings grown on (A) 0.01 µM Cu or (C) 2 µM Cu treated with opening buffer with or without ABA. **(B, D)** Quantification of stomatal aperture in 7-day-old seedlings grown on (B) 0.01 µM Cu or (D) 2 µM Cu treated with opening buffer with or without ABA. Stomata aperture is calculated as the ratio of stomata width to length (mm). The experiment was performed in triplicate, with a total of n = 50 stomata. Levels not connected by the same letter differ significantly (one-way ANOVA, $P<0.01$). Scale bar – 10 µm.

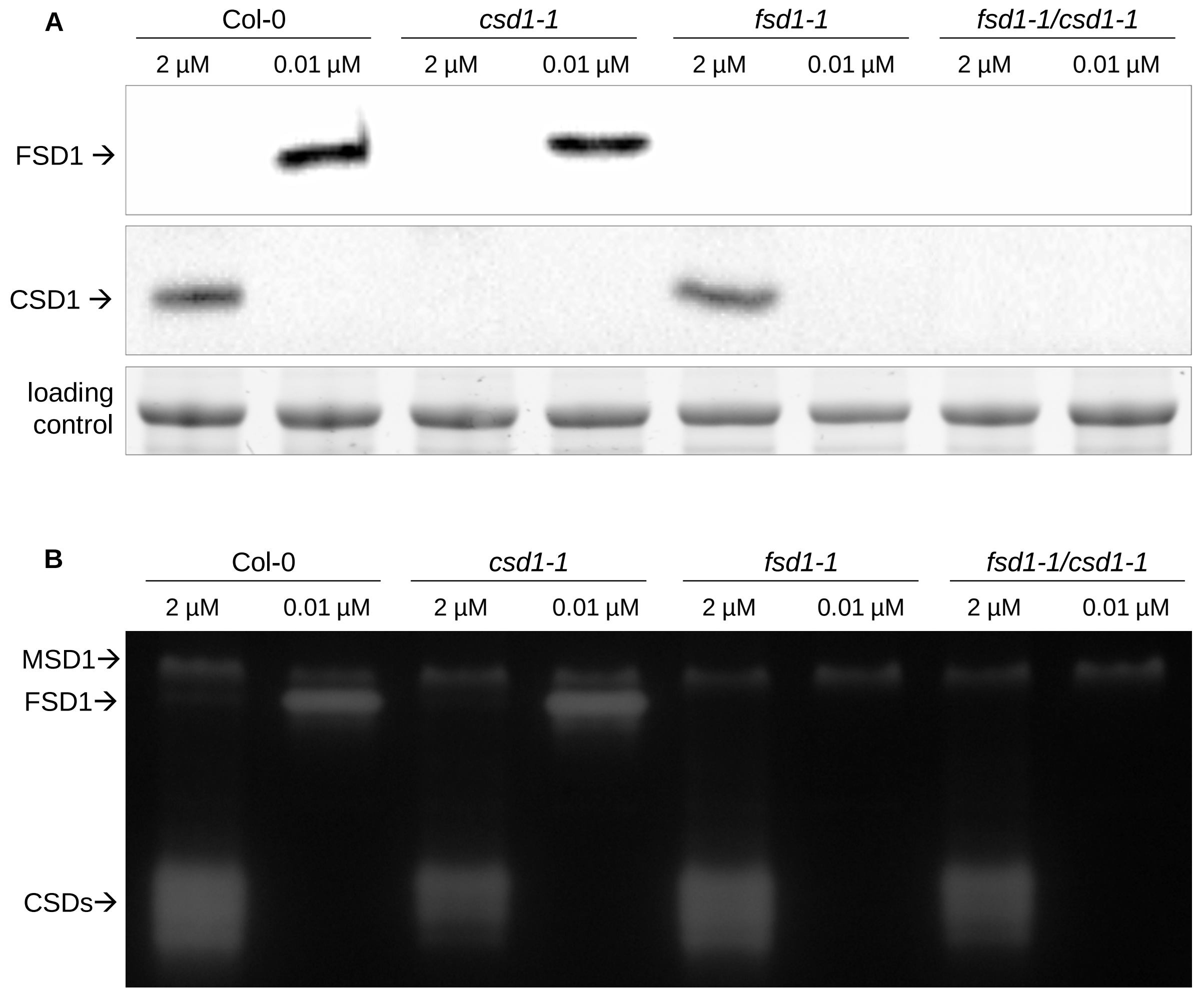


**Supplementary Figure S10. Validation of the double *fsd1/csd1* mutant line. (A)** Immunoblotting analysis of CSD1 and FSD1 abundance in 14-day-old Col-0, *csd1*, and *fsd1* mutants, and a double *fsd1/csd1* mutant line using anti-CSD1 and anti-FSD1 antibodies. **(B)** Visualization of SOD isozymes on native polyacrylamide gel in 14-day-old Col-0, *csd1* and *fsd1* mutants, and a double *fsd1/csd1* mutant line. Seedlings were grown on 0.01 µM Cu (0.01 µM) or 2 µM Cu (2 µM) concentration.

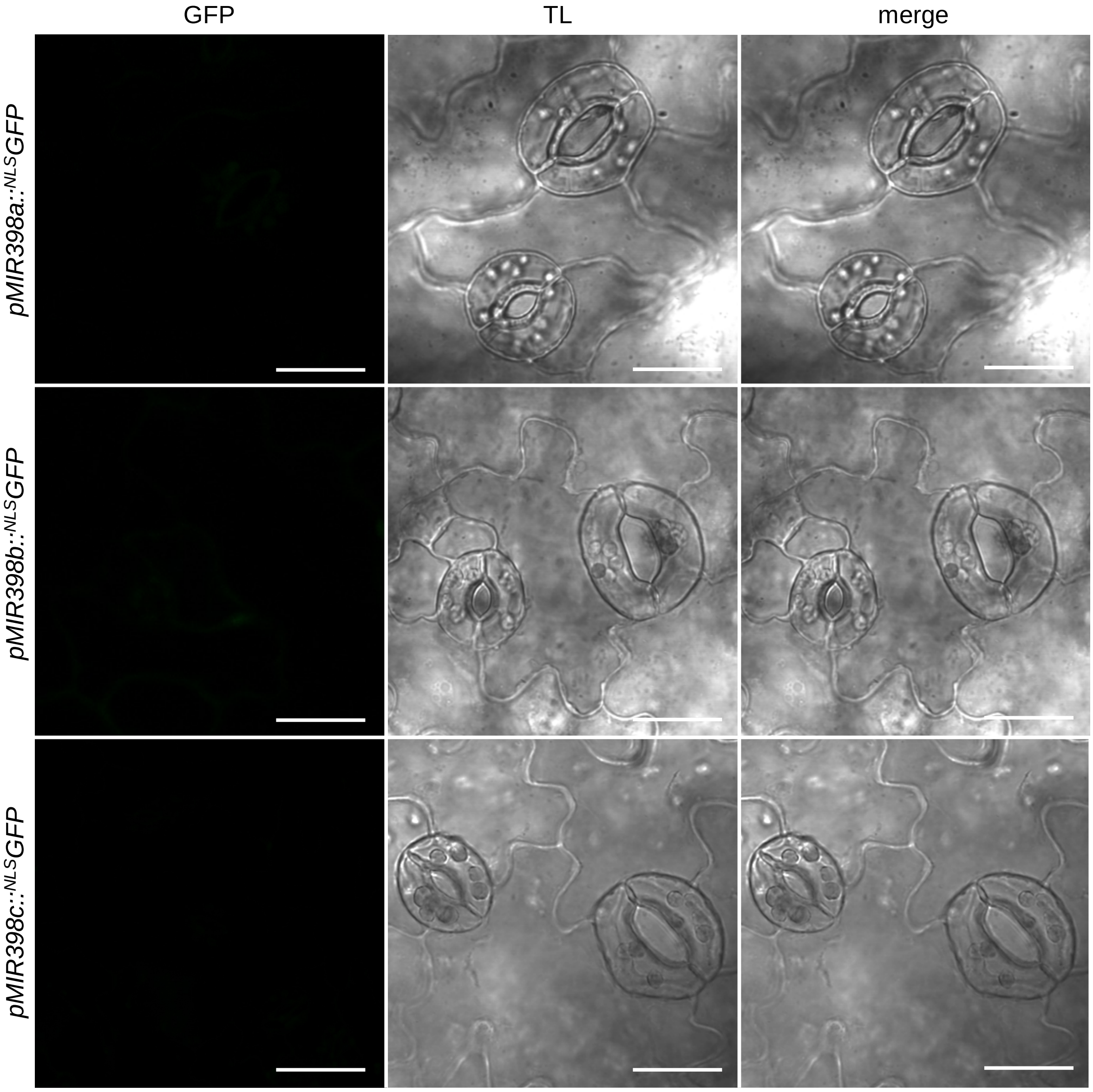


**Supplementary Figure S11. Cu-dependent regulation of *MIR398* promoters.** Representative images of cotyledons of the nuclear-localized GFP lines driven by *MIR398* promoters. Seedlings were grown on 2 μM Cu. TL – transmitted light. Scale bar – 20 μm.

**A**

```
gCSD1        CCCAGCTTTTGGACCTCGTGGGCTATATAGAAAACAAGTAACCAAAGAGAGACGAAGCAAAAACATTCAGAGAGAAAATTCAGCATTTTTGATAGCTCAAGCACTTGATTCTTTCCAAAGGGGTTTCCTGAGGTATGTTTGCTTTC
mRNA CSD1.1  CCCAGCTTTTGGACCTCGTGGGCTATATAGAAAACAAGTAACCAAAGAGAGACGAAGCAAAAACATTCAGAGAGAAAATTCAGCATTTTTGATAGCTCAAGCACTTGATTCTTTCCAAAGGGGTTTCCTGAG--------------
mRNA CSD1.2  CCCAGCTTTTGGACCTCGTGGGCTATATAGAAAACAAGTAACCAAAGAGAGACGAAGCAAAAACATTCAGAGAGAAAATTCAGCATTTTTGATAGCTCAAGCACTTGATTCTTTCCAAAGGGGTTTCCTGAG--------------

gCSD1        CTTTTTGGTTTTAATTTGTTTCAAATCTTAGATCGATTCTTCACTTTGGATTCTTAAATCGATTCTTGTCCTCGTCTGTGCTGATTTCGAACTTTTCCGATTCCAACTCCGCCGAATCTACGTATAGATCTTTGAGGAAATGTTTT
mRNA CSD1.1  --------------------------------------------------------------------------------------------------------------------------------------------------
mRNA CSD1.2  --------------------------------------------------------------------------------------------------------------------------------------------------

gCSD1        TTTCGATTTATGATCGGCGGATCTACCGATTCCTTTGCTTGTTTTCTTTGGATGTATCACACTCTTCTGAAGATGCCTTGACCTTTTTATGTGAACTAGTAAGAGTTATTATGGCTTAGATCGTTGAATATCTGGTATATGTGTTC
mRNA CSD1.1  --------------------------------------------------------------------------------------------------------------------------------------------------
mRNA CSD1.2  --------------------------------------------------------------------------------------------------------------------------------------------------

gCSD1        TTATAATGGGATTAGGATCTATGTTTTGGTGCGTATTAAAGCTTAGTTGAGATTGATGTTTTTGGGTTTTCAAATTCTGTTTCTTGTCTGTTGAGCTAATTTAATGTTGTGTAGAACCATTTGTTTTTTTCTGGATCTTTACTTAT
mRNA CSD1.1  --------------------------------------------------------------------------------------------------------------------------------------------------
mRNA CSD1.2  --------------------------------------------------------------------------------------------------------------------------------------------------

gCSD1        TTGTATATGTATTGGTTTATGTTTATGTTTTTACAGATCACAAAGGCCAAGTAACAATG
mRNA CSD1.1  ------------------------------------ATCACAAAGGCCAAGTAACAATG
mRNA CSD1.2  ---------------------------------------------GCCAAGTAACAATG
```

**B**

*CSD1.2*

FWD1 →

**CCCAGCTTTTGGACC**TCGTGGGCTATATAGAAAACAAGTAACCAAAGAGAGACGAAGCAAAAACATTCAGAGAGAAAATTCAGCATTTTTGATAG

FWD2 →

CTCAAGCACTTGATTCTTTCCAAAGGGGTT**TCCTGAGGCCAAGT**AACAATGGCGAAAGGAGTTGCAGTTTTGAACAGCAGTGAGGGTGTTACGGG

GACTATCTTTTTCACCCAGGAAGGCGATGGTGTGACCACTGTGAGTGGAACAGTTTCTGGCCTTAAGCCTGGTCTTCATGGTTTCCATGTCCATG

CTCTTGGTGACACCACTAACGGTTGCATGTCTACTGGTCCACATTTCAACCCCGATGGTAAAACACACGGTGCCCCTGAGGATGCTAATCGACAT

GCTGGTGATCTAGGAAACATCACTGTTGGAGATGATGGAACTGCCACCTTCACAATCACTGATTGCCAGATTCCTCTTACTGGACCAAACTCTAT

← REV

TGTTGGTAGGGCTGTTGTTGTCCATGCAGACCCTGAT**GACCTCGGAAAGGGA**

*CSD1.1*

FWD1 →

**CCCAGCTTTTGGACC**TCGTGGGCTATATAGAAAACAAGTAACCAAAGAGAGACGAAGCAAAAACATTCAGAGAGAAAATTCAGCATTTTTGATAG

CTCAAGCACTTGATTCTTTCCAAAGGGGTT**TCCTGAG**ATCACAAAG**GCCAAGT**AACAATGGCGAAAGGAGTTGCAGTTTTGAACAGCAGTGAGGG

TGTTACGGGGACTATCTTTTTCACCCAGGAAGGCGATGGTGTGACCACTGTGAGTGGAACAGTTTCTGGCCTTAAGCCTGGTCTTCATGGTTTCC

ATGTCCATGCTCTTGGTGACACCACTAACGGTTGCATGTCTACTGGTCCACATTTCAACCCCGATGGTAAAACACACGGTGCCCCTGAGGATGCT

AATCGACATGCTGGTGATCTAGGAAACATCACTGTTGGAGATGATGGAACTGCCACCTTCACAATCACTGATTGCCAGATTCCTCTTACTGGACC

← REV

AAACTCTATTGTTGGTAGGGCTGTTGTTGTCCATGCAGACCCTGAT**GACCTCGGAAAGGGA**

**Supplementary Figure S12. Alternative splicing of the *CSD1* in the 5`UTR. (A)** Aligned nucleotide sequences of the *CSD1* gene (*gCSD1*), normal splicing variant (*CSD1.1*; NM_100757.4), and alternative splicing variant (*CSD1.2;* NM_001084025.1). Within sequences, the following positions are highlighted: the miR398 binding site in bold; the 5` splice recognition site in blue; the normal 3` splice recognition site in green; the alternative 3` splice recognition site in yellow; the start codon in red. **(B)** Positions of primers in two splice variants. Forward primer (FWD1) is highlighted in yellow, forward primer (FWD2) in blue, annealing only to the *CSD1.2* splice variant; reverse primer (REV) is highlighted in green.

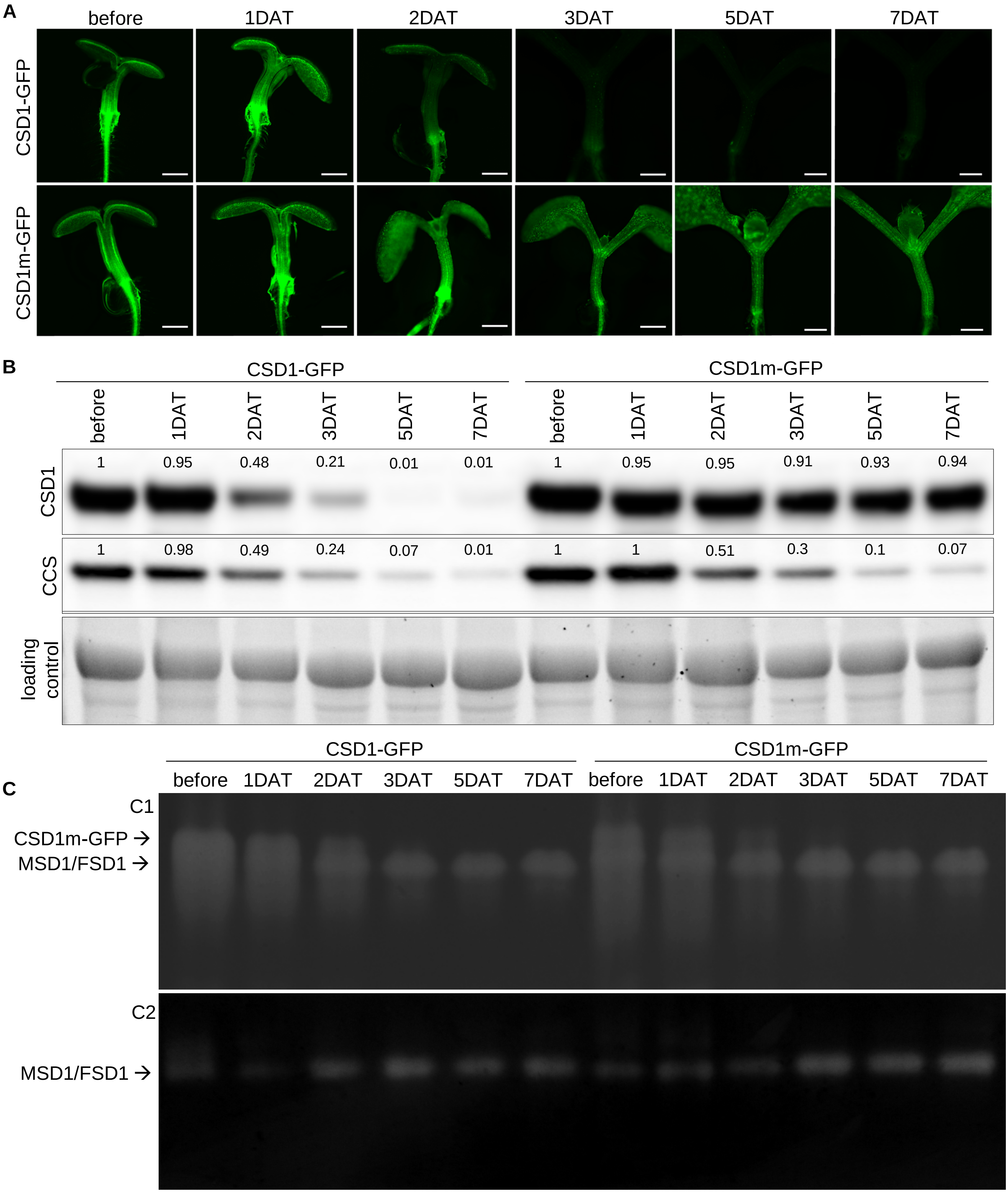


**Supplementary Figure S13. Response to changes in Cu in CSD1-GFP and CSD1m-GFP lines. (A)** Representative images of CSD1-GFP and CSD1m-GFP seedlings before and during seven days after transfer (DAT) from 2 μM to 0.01 μM Cu. **(B-C)** CSD1 and CCS abundance (B) and visualization of SOD isozymes on native polyacrylamide gel (C1) without preincubation and (C2) with preincubation in KCN (which inhibits CSDs' activity) in CSD1-GFP and CSD1m-GFP lines before and during 1- 7 DAT from 2 μM to 0.01 μM Cu. Numbers represent relative changes (nDAT/before) in CSD1 and CCS abundance. Scale bar – 100 μm.

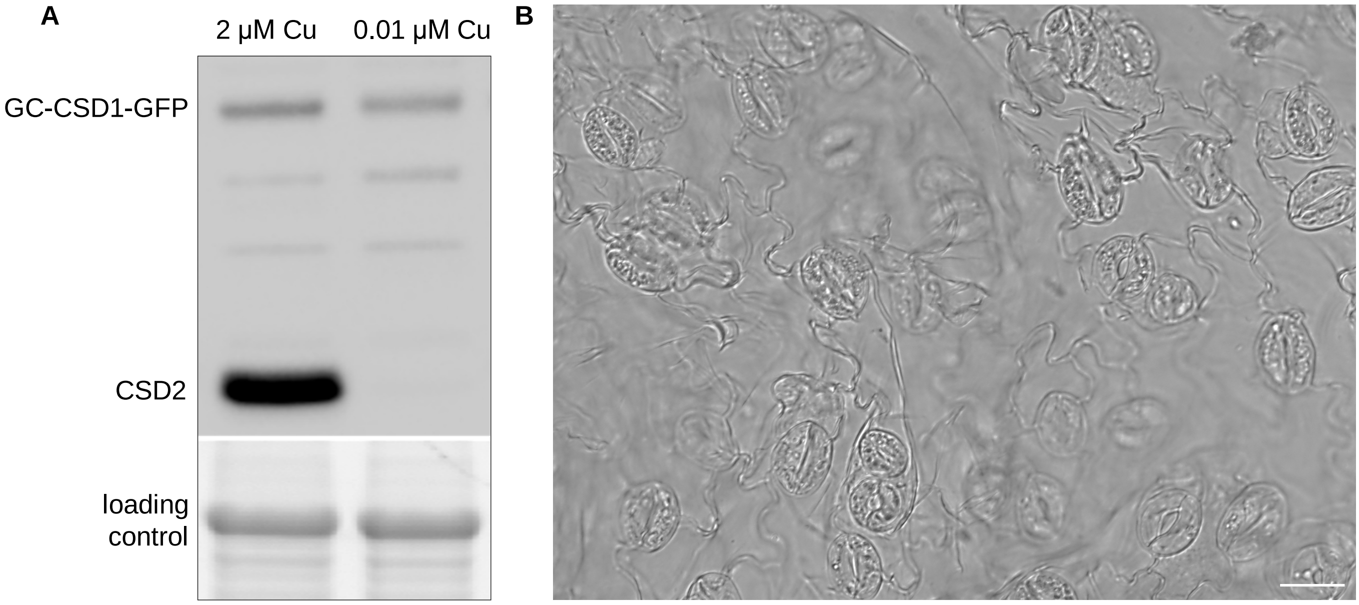


**Supplementary Figure S14. Characteristic of the CSD1 guard cell pool. (A)** The abundance of CSD1 and CSD2 in 14-day-old seedlings of the GC-CSD1-GFP line grown from 2 μM to 0.01 μM Cu. **(B)** Representative image of GC-enriched fraction from 4-week-old seedlings of the GC-CSD1-GFP line grown from 0.01 μM Cu. Scale bar – 20 μm.

```
A  Solyc01g067740.2      CTCTAAGGGTGACCTGAGGTAATTCAATTTTTTTCTTCATCTGTTTTCGATAGTGATTTT   120
   NM_001311084.1        CTCTAAGGGTGACCTGAG------------------------------------------   73
   NM_001247102.2        CTCTAAGGGTGACCTGAG------------------------------------------   63
                                                                  M
   Solyc01g067740.2      ACAGACTCTTGTATCATAATTTTCTAGATCACATACAAAAATG      1123
   NM_001311084.1        ----ACTCTTGTATCATAATTTTCTAGATCACATACAAAAATG      96
   NM_001247102.2        ---------------------------ATCACATACAAAAATG      79

B  Zm00001d029170           GTCGCCTTCCTATCCATCCCTTCCTCGCCGCGGGGGTCGCCTGAGGTATGCTACCTACCA 240
   Zm00001d029170_T003      GTCGCCTTCCTATCCATCCCTTCCTCGCCGCGGGGGTCGCCTGAG--------------- 127
   Zm00001d029170_T004      GTCGCCTTCCTATCCATCCCTTCCTCGCCGCGGGGGTCGCCTGAGGTATGCTACCTACCA 74

   Zm00001d029170           GTTGGATCTCAGATCGGTGGTGCGGTTCCACTTCTAATTTGTTTCCCCCCTTTCCGCTTC 359
   Zm00001d029170_T003      ------------------------------------------------------------ 142
   Zm00001d029170_T004      GTTGGATCTCAGATCGGTG----------------------------------------- 152
                                                                    M
   Zm00001d029170           CCTGTTACTTGTGGACTTGTTTGCAGATCACATAAACAATG      1660
   Zm00001d029170_T003      --------------------------ATCACATAAACAATG      142
   Zm00001d029170_T004      --------------------------ATCACATAAACAATG      167

C  LOC_Os07g46990        CCGCAGGGGTCGCCTGAGGTATGCAGCTTCACCTCCCCCAACTTTCTAGGGTTCTAATCG   180
   LOC_Os07g46990.1      CCGCAGGGGTCGCCTGAG------------------------------------------   138
   LOC_Os07g46990.2      CCGCAGGGGTCGCCTGAG------------------------------------------   60

   LOC_Os07g46990        ACCATCATACTGAGATATATATACCTGTGCACCTTTCGGCTTCTGTGCAGAACACATAGA   720
   LOC_Os07g46990.1      --------------------------------------------------AACACATAGA   148
   LOC_Os07g46990.2      -----------------------------------------------------------A   61
                            M
   LOC_Os07g46990        CAATG  725
   LOC_Os07g46990.1      CAATG  153
   LOC_Os07g46990.2      CAATG  66

D  >Soltu.DM.01G022650                                       M
   CACTCAAGGGTGACCTGAGGTAAT-1121bp-TCTAGATCACATACAAAAATG

   >Medtr7g114240                                           M
   TCTCTGGGGGTTTCCTGAGGTATG-893bp-TTCAGATCACATTGAACAATG
```

**Supplementary Figure S15. Splicing events in the *Cu/ZnSOD1* affecting miR398 sensitivity in various plant species. (A)** Alignment of *Solanum lycopersicum* Cu/Zn superoxide dismutase 1 (Solyc01g067740.2 from the Phytozome database https://phytozome-next.jgi.doe.gov/) and its transcript variants (miR398-insensitive NM_001311084.1; miR398-sensitive NM_001247102.2 from NCBI nucleotide database https://www.ncbi.nlm.nih.gov/). **(B)** Alignment of *Zea mays* Cu/Zn superoxide dismutase 1 (Zm00001d029170) and its transcript variants (miR398-insensitive Zm00001d029170_T003 and miR398-sensitive Zm00001d029170_T004) from the Phytozome database https://phytozome-next.jgi.doe.gov/. **(C)** Alignment of *Oryza sativa* Cu/Zn superoxide dismutase 1 (LOC_Os07g46990) and its transcript variants (miR398-sensitive LOC_Os07g46990.1, miR398-insensitive LOC_Os07g46990.2) from the Phytozome database https://phytozome-next.jgi.doe.gov/. **(D)** Sequence of *Solanum tuberosum* (Soltu.DM.01G022650) and *Medicago truncatula* (Medtr7g114240) Cu/Zn superoxide dismutase 1 from the Phytozome database https://phytozome-next.jgi.doe.gov/. miR398 binding site is highlighted in yellow. Splice sites producing miR398-sensitive variants are highlighted in blue, whereas 5` (for B) or 3` (for A, C) alternative splice sites producing a potentially miR398-insensitive variant are highlighted in green. The start codon (M) is highlighted in bold red. Alignment was done using MEGA software and the Clustal Omega online tool.

**A**

```
CCS  EPLGDLGTLEADKNGEAFYSGKKEKLKV---ADLIGRAVVVYKTDDNKSGPGLTAA 284

CSD1 RHAGDLGNITVGDDGTATFTITDCQIPLTGPNSIVGRAVVVHADPDDLGKGGHELS 133

CSD2 RHAGDLGNINANADGVAETTIVDNQIPLTGPNSVVGRAFVVHELKDDLGKGGHELS 196

CSD3 RHAGDLGNILAGSNGVAEILIKDKHIPLSGQYSILGRAVVVHADPDDLGKGGHKLS 139
        ****                        *** **    *
```

**B**

```
           A  G  D  L  G  N  I
   CSD1 >UGCUGGUGAUCUAGGAAACAUC>
B1       ••••|||||•||••||| ||••
 miR398 >UUCCCCACUGGACUCUU-GUGU>
           A  G  D  L  G  N  I
   CSD2 >UGCGGGUGACCUGGGAAACAUA>
B2       ••|||||||||||•||| ||•|
 miR398 >UUCCCCACUGGACUCUU-GUGU>

   CSD1 >AAGGGGUUUCCUGAGAUCACA>
B3       |||||||••|||||||•||||
 miR398 >UUCCCCACUGGACUCUUGUGU>
```

**Supplementary Figure S16. Localization of the miR398 binding site in *Arabidopsis* Cu/ZnSOD (CSD1, CSD2, CSD3) and COPPER CHAPERONE FOR SUPEROXIDE DISMUTASE (CCS). (A)** Alignment of CSD1, CSD2, CSD3, and CCS protein sequences. miR398 binding site is highlighted in yellow. Glycine (G) incorporated in the active site is highlighted in bold and underlined. Stars indicate conservative amino acids across all aligned sequences. **(B)** Scheme of *miR398* binding to the (B1) *CSD1* mRNA in the coding sequence (highlighted in grey in A); (B2) *CSD2* mRNA in the coding sequence (highlighted in yellow in A); (B3) *CSD1* mRNA in the 5`UTR (shown in S14). Letters above represent the amino acid coded by a triplet of nucleotides. The dots represent the disruption of the interaction between individual mRNA nucleotides and the mature miR398. The dashes indicate complementarity within the interaction framework.
