## Supplementary table 1 for "Stomatal movement in *Arabidopsis* is driven by guard cell-localized and copper-insensitive CSD1 splice variant"

**Supplementary Table 1.** List of used primers.

| Primer name | Primer sequence | Use |
|---|---|---|
| LBb1.3 | ATTTTGCCGATTTCGGAAC | PCR |
| SALK_109389-LP | TCTTCTGAAGATGCCTTGACC | PCR |
| SALK_029455-LP | TTTGTTTGGTCTCCCAACAAC | PCR |
| SALK_024857-LP | ATGAACCCCGAGTTACCAGAG | PCR |
| SALK_109389-RP | GTCATTACCCTTTCCGAGGTC | PCR |
| SALK_029455-RP | GTTGAAAGCGAGGAGATC | PCR |
| SALK_024857-RP | TTGCAGTTTTGAACAGCAGTG | PCR |
| C-terminal fusion: Promoter + gene_FWD | GGGGACAACTTTGTATAGAAAAGTTGCGGCTTCCAAATGGTTGAATCAG | PCR |
| C-terminal fusion: Promoter + gene_REV | GGGGACTGCTTTTTTGTACAAACTTGCGCCCTGGAGACCAATGATG | PCR |
| C-terminal fusion: 3´UTR_FWD | GGGGACAGCTTTCTTGTACAAAGTGGCGAGCTGCTACGTTTCCAAAG | PCR |
| C-terminal fusion: 3´UTR_REV | GGGGACAACTTTGTATAATAAAGTTGCCTTGTATCTCCATTTTTGTCC | PCR |
| pDONR.1_CSD1m_FWD | CTTGATTCTTTCCAAAAGCTTTTCGCGAGGTATGTTT GCTTTCCTTTTTGGTTTTAATTTG | PCR |
| pDONR.1_CSD1m_REV | GAAAGCAAACATACCTCGCGAAAAGCTTTTGGAAAG AATCAAGTGCTTGAGC | PCR |
| pDONR.2_CSD1m_FWD | GCGAGACGAAATACGCGATCGCTGTTAAAAG | PCR |
| pDONR.2_CSD1m_REV | ACAGCGATCGCGTATTTCGTCTCGCTCAGGC | PCR |
| pENTR.1_MIRa_FWD | CAACAGGAGGGACATGCAGCCTTCTCTTAAAC | PCR |
| pENTR.1_MIRa_REV | GCATCGACTACGCCAGCCTGCTTTTTTG | PCR |
| pENTR.2_MIRa_FWD | CAGGCTGGCGTAGTCGATGCGGGATGTTG | PCR |
| pENTR.2_MIRa_REV | GCTGCATGTCCCTCCTGTTGTTCTTCTCTTC | PCR |
| pENTR.1_MIRb_FWD | TTCAGTCGACCTGAATGCAATCATACAAAGAAG | PCR |
| pENTR.1_MIRb_REV | AAGGCTGCATCTACCTTCATGATATATTGATATCTTCTTG | PCR |
| pENTR.2_MIRb_FWD | ATGAAGGTAGATGCAGCCTTCTCTTAAACG | PCR |
| pENTR.2_MIRb_REV | TTGCATTCAGGTCGACTGAATTGGTTCCG | PCR |
| pENTR.1_MIRc_FWD | GTTAAGGTAGGACATGCAGCCTTCTCTTAAAC | PCR |
| pENTR.1_MIRc_REV | ACCTCTGTCTCGCCAGCCTGCTTTTTTG | PCR |
| pENTR.2_MIRc_FWD | CAGGCTGGCGAGACAGAGGTATCATTCACC | PCR |
| pENTR.2_MIRc_REV | GCTGCATGTCCTACCTTAACAATATTTATCTATCTTCTAAAC | PCR |
| pDEST.1_CSD1m_FWD | CCAAGCTATCGTATAGAAAAGTTGCGGCTTC | PCR |
| pDEST.1_CSD1m_REV | TGCTCACCATGCAGCCTGCTTTTTTGTAC | PCR |
| pDEST.2_CSD1m_FWD | AGCAGGCTGCATGGTGAGCAAGGGCGAG | PCR |
| pDEST.2_CSD1m_REV | GTGGCTGGCTCGAAGATACCTGCAAGAATG | PCR |
| pDEST.3_CSD1m_FWD | GGTATCTTCGAGCCAGCCACGATCGACATTG | PCR |
| pDEST.3_CSD1m_REV | TTTTCTATACGATAGCTTGGCGTAATCATGGTC | PCR |
| pDEST.1_GC_FWD | CCAAGCTATCTGAATTTATAAGTTTTCAACACCG | PCR |
| pDEST.1_GC_REV | CTTTCGCCATTAGTGTGTGAAAATAGTACTTGTG | PCR |
| pDEST.2_GC_FWD | TCACACACTAATGGCGAAAGGAGTTGCAG | PCR |
| pDEST.2_GC_REV | GTGGCTGGCTCGAAGATACCTGCAAGAATG | PCR |
| pDEST.3_GC_FWD | GGTATCTTCGAGCCAGCCACGATCGACATTG | PCR |
| pDEST.3_GC_REV | TATAAATTCAGATAGCTTGGCGTAATCATGGTC | PCR |
| FWD1_splice | CCCAGCTTTTGGACC | PCR |
| FWD2_splice | TCCTGAGGCCAAGT | PCR |
| REV1_splice | TCCCTTTCCGAGGTC | PCR |
